# Weakly supervised identification and generation of adaptive immune receptor sequences associated with immune disease status

**DOI:** 10.1101/2023.09.24.558823

**Authors:** Andrei Slabodkin, Ludvig M. Sollid, Geir Kjetil Sandve, Philippe A. Robert, Victor Greiff

## Abstract

Adaptive immune receptor (AIR) repertoires carry immune signals as sequence motif imprints of past and present encounters with antigen (immune status). Machine learning (ML)-based identification and generation of antigen-specific immune receptors is potentially immense value for public health. The ideal training data for such ML tasks would be AIR datasets, where each sequence is labeled with its cognate antigen. However, given current technological constraints, sequence-labeled datasets are scarce, contrasted by an abundance of repertoire-labeled ones – AIR repertoire datasets where only the repertoire dataset, but not the individual AIRs, are labeled. Therefore, an unmet need exists for an ML approach that enables predictive identification and generation of disease-specific novel AIR sequences using exclusively repertoire-level immune status information. To address this need, we developed AIRRTM, an end-to-end generative model using an encoder-decoder architecture and Topic Modeling (TM) that requires exclusively repertoire-labeled AIR sequencing data as input. We validated AIRRTM’s capacity to identify and generate novel disease-associated receptors on several ground truth synthetic datasets of increasingly complex immune signals and experimental data. AIRRTM broadens the discovery space for immunotherapeutics by enabling the exploitation of large-scale and broadly available immune repertoire data previously deemed largely unsuitable for this task.

## Introduction

Adaptive immune receptor (AIR) repertoires are responsible for the specific recognition and neutralization of pathogens (Rappazzo et al. 2023) and are of importance for the diagnostics and treatment of disease and infection (Greiff et al. 2020). So far, the immense intra- and inter-individual diversity of immune receptors renders it particularly challenging to learn the association between immune receptor and immune status (Briney et al. 2019; Greiff et al. 2017; Shemesh et al. 2021; Gidoni et al. 2019; Emerson et al. 2017). By immune status, we mean any condition affecting the individual’s AIR repertoire. For example, it can be an ongoing immune response to infection, immune competence to fight infection (i.e., protection provided by vaccines), the presence of a tumor, or different stages of immune-related diseases. Even more challenging is the task of generating novel sequences that satisfy certain requirements, such as specificity to a given antigen or antibody developability (Akbar et al. 2022a, 2022b; Saka et al. 2021; Amimeur et al. 2020; Hie et al. 2023).

A typical AIR dataset is either a set of repertoires (i.e., sets of AIR sequences) of subjects labeled with several immune states (i.e., HIV-positive and HIV-negative or celiac-disease-affected and non-affected subjects), or a list of AIR sequences labeled with binding to one or more target antigens. High-throughput AIR repertoire-sequencing (AIRR-seq) (Weinstein et al. 2009) allows to generate amounts of experimental data large enough to cover the immune receptor diversity: tens of millions of immune receptors can be sequenced at a relatively low expense, and billions of sequences are already deposited in public databases (Kovaltsuk et al. 2020; Corrie et al. 2018). At the same time, experimental methods for identifying and annotating antigen-specific sequences (Stiegler et al. 2001; Wu et al. 2010; Walker et al. 2011; Setliff et al. 2019), while progressing rapidly, still lag behind antigen-agnostic bulk AIRR-sequencing by several orders of magnitude, both in terms of cost-per-sequence and number of publicly available immune receptor sequences annotated with antigen specificity (Raybould et al. 2020; Swindells et al. 2017; Ferdous and Martin 2018; Rappazzo et al. 2023). Thus, a method for identifying individual immune receptors that determine the immune status of the repertoire (as well as generating novel sequences with the same property) may be of great benefit, given the large amount of publicly available bulk AIRR-sequencing data with associated immune status metadata (Olsen et al. 2022a; Corrie et al. 2018). Whether it is possible to develop such a method and whether the signal-to-noise ratio in the currently available datasets is sufficient are the main research questions of this work.

Adaptive immune receptor repertoire labels (e.g., immune status of a given individual) are usually available even for bulk AIRR-seq (Corrie et al. 2018; Olsen et al. 2022a). In this study, we formalize learning of the association between immune receptor and immune status from AIR repertoires as two weakly-supervised ML tasks (Zhou 2018): identification of signal-positive immune receptor sequences (signal sequence identification) from repertoires knowing the immune status of each repertoire, and signal-associated sequence generation, i.e., generation of new immune receptor sequences associated with immune status. These two tasks may have direct implications for diagnostics and therapeutics of infectious and autoimmune diseases: *in silico* signal sequence identification would allow finding and/or designing candidate sequences of antigen-binding immune receptors from the repertoire-labeled AIRR data alone.

Emerson et al. 2017; Widrich et al. 2020; Gao et al. 2023, and Chen et al. 2023 developed methods for signal sequence identification using only repertoire labels but not sequence labels. Daberdaku and Ferrari 2019, Akbar et al. 2021, Lu et al. 2020 showed that signal sequence identification can be solved given a sufficiently large amount of data with sequence labels. Eguchi et al. 2020; Friedensohn et al. 2020; Akbar et al. 2022b used generative models trained on labeled sequences to produce novel label-specific sequences. Choi 2022, Ruffolo et al. 2021, Olsen et al. 2022b trained larger generative models on individual antibody sequences. Marcou et al. 2018 used an explicit Bayesian probabilistic model (Olson and Matsen 2018), to generate AIR repertoires that mimic experimental ones regardless of any sequence labels, while Davidsen et al. 2019 and Isacchini et al. 2021 used deep generative models for the same goal. Most importantly, Pradier et al. 2023 designed a generative model AIRIVA that can learn a low-dimensional interpretable representation of TCR repertoires with respect to the repertoire labels.

So far, however, no method solves both the identification of signal sequences and the generation of novel signal sequences using only [relatively inexpensive and readily available] repertoire labels but not [expensive and scarce] sequence labels. Of note, the previously reported AIRIVA model by Pradier et al. (Pradier et al. 2023) identifies signal sequences and generates diverse repertoires containing these sequences. Still, generating novel sequences is out of its scope (see Methods 2 for details). Here, we present such a method: Adaptive Immune Receptor Repertoires Topic Modeling (AIRRTM). On synthetic and semi-experimental data, AIRRTM was able to identify signal-positive sequences in the data robustly and to generate novel signal-positive sequences via learning from repertoire-label-only data. Another challenge is that there may be only a few signal-positive sequences in the data (low *witness rate*). We show that AIRRTM produces consistent results even for a witness rate as low as 1 signal-positive sequence per 10000, which is in line with witness rates observed in experimental data (Abbott et al. 2018). In the absence of ground truth labels for individual sequences, AIRRTM performed classification of unseen repertoires on par with the previous state-of-the-art methods applied to the same data.

## Results

### An ML architecture to infer and generate signal-positive sequences from immune status-labeled repertoires

We start by defining the machine learning task and then applying it to bulk AIRR-seq data (Fig. 1A). Given a set of (experimental or synthetic) AIR repertoires, each repertoire is a collection of AIR (here: CDR3) amino acid sequences. The repertoires are immune-status labeled (Fig. 1B): some repertoires are signal-positive (e.g., infected or diseased), and the rest are signal-negative (e.g., healthy controls). We assume that signal-positive repertoires contain signal-positive AIR sequences (Fig. 1C): those sequences satisfy a certain condition (for example, autoreactive sequences when the immune status is an autoimmune disease). We employ a general definition of signal-positive sequence without specifying a type of sequence motif or explicit sequence rules that render a sequence signal-positive since our ML formulation could work with any set of sequences being signal-positive. In this study, all signal-positive repertoires are assumed to have the same fraction of signal-positive sequences (witness rate). The repertoire labels are available to the ML method, but the information on which sequences are signal-positive is not. The goal is to (i) identify signal-positive sequences and to (ii) generate novel sequences that are signal-positive; in settings with ground-truth definition of signal-positive sequences, the generated sequences should meet the same definition. We built our approach upon Topic Modeling (TM), a technique (Hofmann 1999; Blei et al. 2001) widely used in the neighboring field of Natural Language Processing. TM is a set of methods for identifying interpretable patterns (or “topics”) in a collection of texts (see Methods 3 for details). These methods usually represent a text as a bag-of-words where the word order in the text does not matter. This assumption, obviously incorrect for natural language texts, is much closer to reality in the case of (naïve) AIR repertoires (Vu et al. 2022, 2023). However, there are significant differences between AIR sequences and natural language words and phrases: the diversity of AIR sequences is substantially higher – 10^6^ for English words versus 10^12^ for naïve antibody repertoire (Briney et al. 2019) and the sequence space is more uniform (in terms of distance between sequences/words). Hence, character-level information is more important for modeling. To adapt Topic Modeling to AIRR-seq data, we developed AIRRTM, combining several approaches: Additive regularization of Topic Models (Vorontsov and Potapenko 2015), Embedded Topic Model (Dieng et al. 2020) and Sequence VAE (Chung et al. 2015), which allowed us to connect the latent space learned by the sequence variational autoencoder to the topics in repertoires (see Methods 3 and Fig. 2E for details).

**Figure 1:**
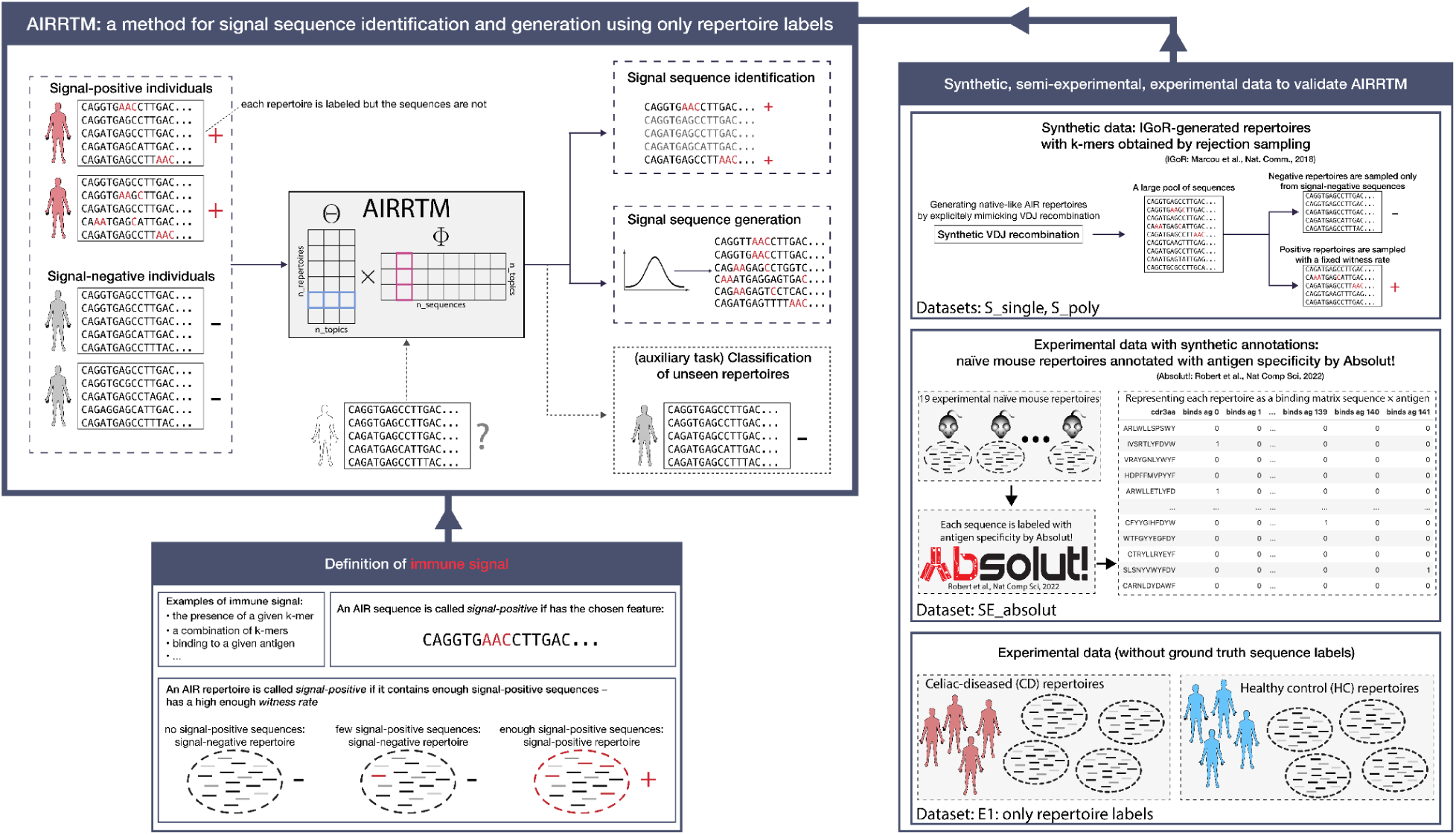
Weakly supervised identification and generation of signal-associated adaptive immune receptor sequences using repertoire labels but not individual sequence labels. AIRRTM allows approaching both signal sequence identification and signal sequence generation using only repertoire-labeled AIRR sequencing data. We provide definitions of immune signal for AIR repertoires and AIR sequences. We created datasets of increasing complexity and naturalness to validate AIRRTM: we used IGoR (Marcou et al. 2018) to generate synthetic AIR repertoires and obtained sequences containing specific sequences with rejection sampling. We used Absolut! (Robert et al. 2022) to annotate experimental naïve mouse repertoires from (Greiff et al. 2017) with specificity to 142 antigens. We also applied AIRRTM to experimental data without ground truth individual sequence labels.

**Figure 2:**
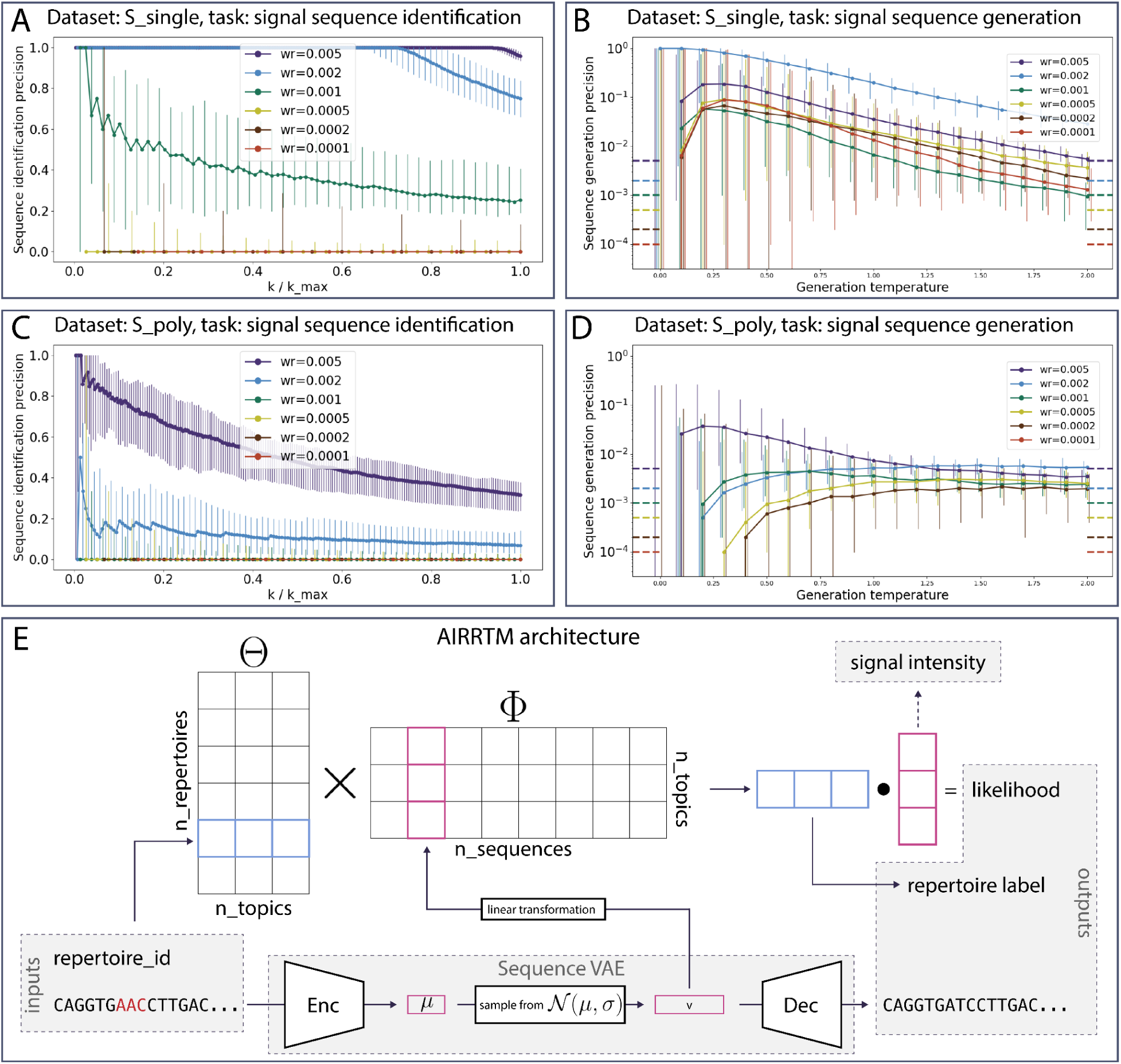
AIRRTM identifies signal-positive sequences and generates novel signal-positive sequences for single-signal and multi-signal ground truth synthetic datasets using only repertoire-level labeled data as training input. (A) Signal sequence identification precision for the S_single dataset (50 signal positive synthetic repertoires containing sequences with amino acid 3-mer DPM, 50 synthetic signal-negative repertoires, each repertoire contains 80000 sequences in total – see Methods 1 for details): all sequences in a repertoire are ranked by their signal intensity as predicted by AIRRTM. We report the fraction of correctly identified signal-positive sequences (median across repertoires, error bars show the total range from minimum to maximum across repertoires) among top `k` predictions (`k` in `1..k_max`), `k_max = witness_rate·sample_size`. The control identification precision value equals the witness rate and thus cannot be shown in the plot, as it is too low. “wr” stands for witness rate. (B) Signal sequence generation precision for the dataset S_single. X axis: generation temperature. Left Y axis, solid lines: fraction of sequences that bear the signal among 100000 generated sequences (median across repertoires, error bars show the total range from minimum to maximum across repertoires), dotted lines correspond to the random baseline, i.e., witness rate). (C) Same as A but for the S_poly dataset (synthetic repertoires where the signal is one of the four different amino acid 3-mers: QGD, IKL, ENQ, SPF; in different proportions). (D) Same as B but for the S_poly dataset. (E) Architecture of the AIRRTM model: (i) a pair (AIR sequence, repertoire_id) is taken as input, (ii) the sequence is encoded to obtain a latent space representation (encoding), (iii) a linear operation with sigmoid transforms the obtained encoding vector into sequence topic probabilities, constituting a column of the `topic×sequence` matrix Φ, (iv) using a lookup by repertoire_id, a repertoire topic proportion vector – a row of the `repertoire×topic` matrix Θ – is obtained, (v) the dot product of sequence topic probabilities and repertoire topic proportions is computed as likelihood of observing the given sequence in the given repertoire – output 1, used to compute the TM loss component, (vi) a repertoire label is predicted by logistic regression on the repertoire topic proportion vector – output 2, used label prediction loss component, (vii) the sequence latent vector is decoded – output 3, used to compute the reconstruction loss component (see Methods 3 for details).

### AIRRTM identifies signal-positive sequences and generates novel signal-positive sequences from single-signal ground truth synthetic datasets

We first assessed whether AIRRTM can infer simple types of immune signals based on synthetic AIR repertoire datasets. Here, ground-truth signals were manually added to some repertoires, which are labeled with a corresponding immune status – a fully controlled synthetic setting (Sandve and Greiff 2022). In each dataset, the ground truth individual sequence labels are available, which allows us to evaluate the performance of the designed model. We first generated a dataset of 100 repertoires, each containing 80000 CDRH3 sequences (**`S_single`**, see Methods 1 for details). The repertoires were generated with realistic V(D)J recombined BCR sequences using the IGoR model (Marcou et al. 2018) using identical recombination parameters for each repertoire (Slabodkin et al. 2021). Then, a single amino acid 3-mer was chosen to represent the signal – representing immune signals as k-mers (i) has been widely adopted in the field (Akbar et al. 2021; Kanduri et al. 2021; Ostmeyer et al. 2019) and (ii) has been shown to reflect the natural binding patterns (Akbar et al. 2021; Ostmeyer et al. 2019; Shrock et al. 2023). 50 repertoires were generated without any sequences containing the signal (signal-negative repertoires), and the other 50 were generated with a fixed fraction of sequences containing the signal (witness rate). We generated the 100 repertoires with multiple different values of witness rate separately, thus generating multiple different versions of the dataset. The minimum (0.01%) and maximum (0.5%) witness rates were chosen so that the performance of baseline repertoire classification spans the whole range from random to perfect, as previously determined by Kanduri et al. 2021. Since the background repertoires were generated with IGoR (Marcou et al. 2018) and signal-positive sequences were obtained with rejection sampling (see Methods 1), we did not specifically control where the signal k-mer was located within the sequence. In this dataset, it resulted in the k-mer naturally occurring at the same position (the 3rd amino acid from the 5’ end) in the majority of signal-positive sequences (Fig. S3A), which rendered the signal less complex.

AIRRTM was trained on three objectives simultaneously: (i) sequence encoding-decoding, (ii) repertoire label prediction based on the topic proportions of the repertoire, and (iii) prediction of whether a sequence originated from a repertoire based on the predicted topic probabilities for the sequence and the topic proportions of the repertoire (see Methods 3 and Fig. 2E for details). The learned representations of sequences – predicted topic probabilities – were used to calculate signal intensity that indicates how much the model is confident that a sequence is signal-positive (see Methods 3).

To evaluate the model performance for signal sequence identification, we ranked all sequences from a signal-positive repertoire *unseen by the model* (test set) by the signal strength predicted by the trained AIRRTM model. For each `k` ∊ `1..k_max`, we calculated the sequence identification precision (Fig. 2A): `n_pos/k`, where `n_pos` is the number of signal-positive sequences among the `k` top-ranked, and `k_max` is the total number of signal-positive sequences in the dataset – which, depending on the witness rate, varies from 400 (wr=0.5%) to 8 (wr=0.01%). For the described dataset, AIRRTM showed sufficient signal sequence identification performance (Fig. 2A). For higher witness rates (0.1%–0.5%), the first several top predictions were perfectly correct, then the precision scaled down to 0.9–0.5; for lower witness rates, the precision was close to zero. However, the predicted signal intensity (used to rank the sequences for signal identification) distribution for signal-negative and signal-positive sequences for all witness rates (Fig. S1, left column) shows that even for the lowest witness rate AIRRTM successfully captures the signal, as the distribution of predicted signal intensities for signal-positive sequences is skewed compared to the signal-negative one.

For signal sequence generation, we (i) chose `k_max` identified sequences, (ii) computed their latent space representation vectors, (iii) computed the mean and standard deviation, of these `k_max` vectors, and (iv) sampled 100000 vectors from a normal distribution with mean μ and covariance matrix Diag((*σ*·t)^2^), where `t` is the generation temperature, and decoded these vectors into sequences. Then, we measured how many of these generated sequences are signal-positive and how many of them are novel and unique (Fig. 2B). We repeated the process for multiple values of `t` ∊ `[0,2]`: higher generation temperature yielded a more diverse set of sequences, but typically lower precision (Fig. S4). Here, even for the lower witness rates, AIRRTM achieves single-digit percent signal sequence generation precision, which substantially supersedes the random performance, i.e., the witness rate (see Discussion for further interpretation of precision values and their sufficiency for experimental application). Therefore, we showed that AIRRTM consistently identifies signal sequences and generates novel signal sequences in single-signal synthetic data.

### AIRRTM identifies signal-positive sequences and generates novel signal-positive sequences from multi-signal ground truth synthetic datasets

To validate AIRRTM in a more challenging environment, we created a synthetic dataset containing four different immune signals (S_poly, see Methods 1 for details): same as for the S_single dataset, we generated 100 repertoires with identical VDJ recombination parameters. Here, we chose four different 3-mers with different natural occurrence rates (Fig. S3, see Methods 1 for details) to represent the signal. Again, as with the S_single dataset, 50 repertoires were generated without any sequences containing those 3-mers, and the other 50 were generated with a fixed fraction of sequences containing any of the chosen 3-mers. For each repertoire separately, we sampled signal proportions from the uniform(0,1) distribution. and rescaled them so that each signal-positive repertoire normally contained all four 3-mers, and the total frequency of signal-positive sequences was equal to the witness rate. This way, the signal was diluted between the 3-mers, which made the task more complex. Another complexity when compared to the S_single dataset is the positions at which these 3-mers naturally occur (Fig. S3). While the 3-mer in the S_single dataset mostly starts at the same position close to the beginning of the sequence, the four 3-mers from the S_poly dataset were distributed more evenly.

For the S_poly dataset, AIRRTM showed worse sequence identification precision than for S_single. However, it still was able to correctly identify signal-positive sequences for the three higher witness rates (Fig. 2C). Although the predicted signal intensity levels varied across the signal 3-mers – for higher witness rates, sequences containing ENQ had on average 2.1 times higher predicted signal intensity than sequences containing SPF – the model was able to capture all four components of the signal (Fig. S2, left column). However, the model cannot reliably separate different components of the signal (Fig. S2, right column), which may be the reason behind the drop in signal sequence generation performance (compared to the S_single dataset). We thus showed that AIRRTM consistently identifies signal sequences in multi-signal synthetic data and consistently generates novel signal sequences, although possibly not for all components of the signal.

### AIRRTM enables signal sequence identification and signal sequence generation for native murine data with complex synthetic signal sequence annotations

To test our model on even more complex signals than explored above, we used Absolut! (Robert et al. 2021). Absolut! is a software package for the generation of lattice-based 3D-antibody-antigen binding structures with ground-truth access to conformational paratope, epitope, and affinity. We used it to annotate CDRH3 sequences from nineteen naïve mouse repertoires (Greiff et al. 2017) with Absolut!-defined specificity to 140 antigens (SE_absolut dataset, Fig. 3A). Of note, we do not claim that Absolut!-defined bindings recapitulate experimental ones: the signal defined by these bindings is complex and non-trivial, and we can use it as an estimate of the experimental complexity of immune signals representing antigen specificity. In other words, this dataset presents a problem of similar difficulty to the experimental data with non-trivial signal-positive sequence patterns.

**Figure 3:**
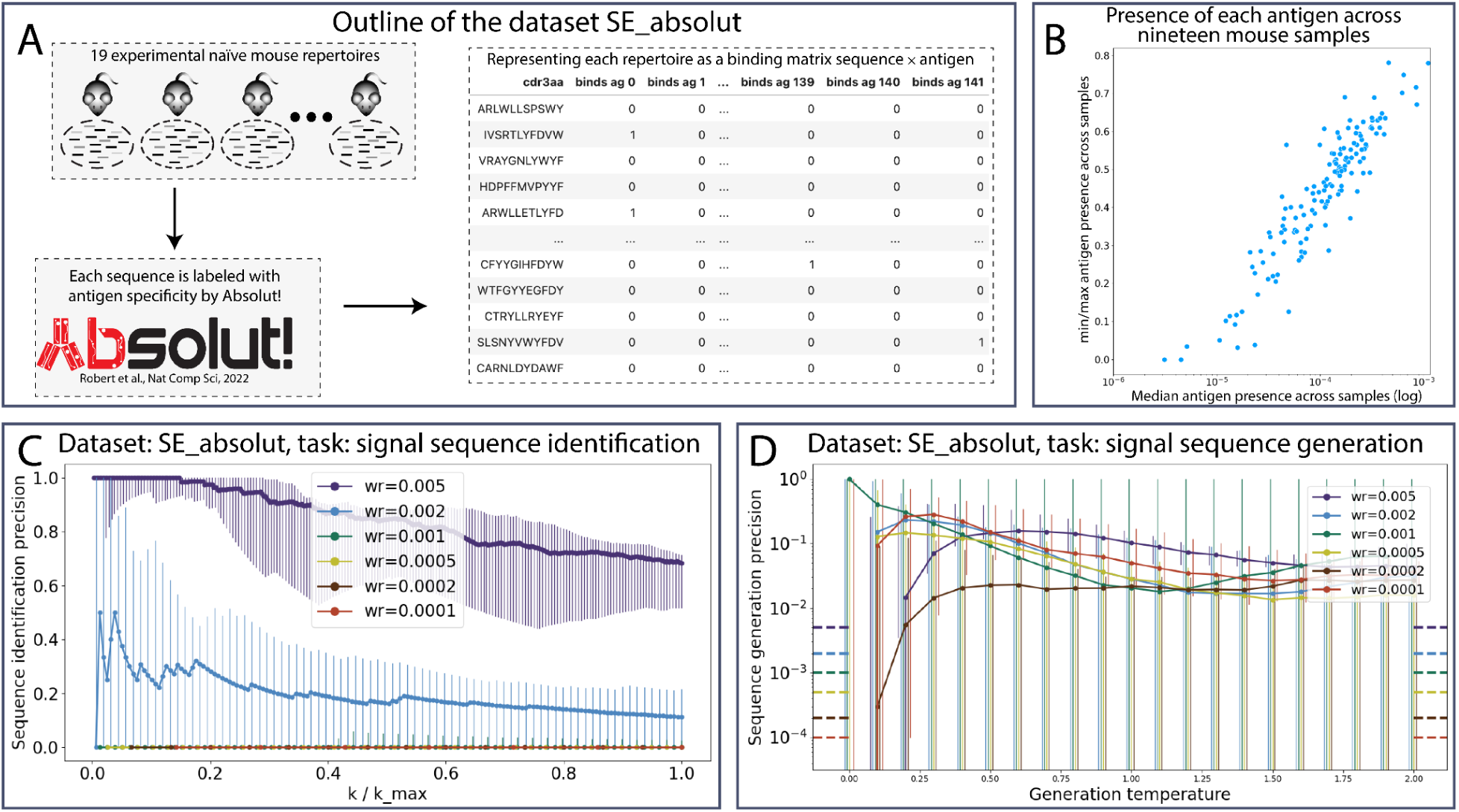
AIRRTM identifies signal-positive sequences and generates novel signal-positive sequences for experimental data with complex synthetic signal sequence annotations using only repertoire labels. (A) Natural mouse repertoires are annotated with antigen specificity by the Absolut! framework (Robert et al. 2022, see Methods 1 for details). (B) Antigen binders are distributed similarly across 19 experimental repertoires: minimal-to-maximum ratio of antigen presence across repertoires (Y axis) is lower than 0.2 only for antigens with very low presence (X axis). Witness rate for binders to different antigens across samples of the SE_absolut dataset. (C) AIRRTM signal sequence identification precision for the SE_absolut dataset (18 natural mouse repertoires, where the signal is Absolut!-defined binding): fraction of correctly identified signal-positive sequences (median across repertoires, error bars show the total range from minimum to maximum across repertoires) among top `k` sequences with highest predicted signal intensity (`k` in `1..k_max`), `k_max = witness_rate·sample_size`. The control identification precision value equals the witness rate and thus cannot be shown in the plot, as it is too low. (D) Signal sequence generation precision for the dataset SE_absolut. X-axis: generation temperature. Y axis, solid lines: fraction of sequences that bear the signal among 10000 generated sequences (median across repertoires, error bars show the total range from minimum to maximum across repertoires), dotted lines correspond to the control value, i.e., witness rate).

To validate AIRRTM, we need the repertoire labels (the presence of an Absolut!-defined binders to a given antigen in the repertoires) to be different between repertoires. To check this, we calculated the presence of sequences binding each antigen across (antigen presence) the nineteen repertoires (Fig. 3B): in most cases, the frequencies of binders were similar across repertoires – thus, these repertoires cannot be used as a dataset for the ML tasks studied here (signal sequence identification and generation), since every repertoire is positive for each label, and naïve repertoires could have a pre-existing coverage of many possible antigens, while immune signals are typically derived from non-homeostatic events.

To adapt the dataset for these tasks, we created the SE_Absolut dataset with modified positive or negative repertoires by increasing or decreasing their amount of signal (here, we did this for a single antigen, PDB ID: 1NSN). For a given antigen, we selected nine repertoires with the lowest frequencies of antigen-binding sequences as negative repertoires, and removed all such binders from them. Then, for building positive repertoires, we decreased the amount of antigen binding to an antigen to reach a given witness rate – those repertoires are considered positive. Therefore, the design of positive and negative repertoires follows the same setting as in S_single and S_poly, only with a different signal. This signal, however, is much more complex than a combination of k-mers, as it encompasses the 3D structural relationship between antibodies and the target antigen. In addition to that, the repertoires as sets of sequences contain all the complexity of experimental data in terms of sequence diversity.

For this scenario, AIRRTM showed performance similar to the S_single dataset: for the higher witness rates of antigen-binding sequences, the signal sequence identification precision ranged from above ten percent to perfect (Fig. 3C), and even for the witness rate 0.05%, although there were few or none signal-positive sequences among the top-ranked ones, AIRRTM was still able to capture the signal, as shown by the predicted signal intensity for signal-positive (binders) and signal-negative (non-binders) sequences (Fig. S5, left column). At the same time, for almost all witness rates, AIRRTM yielded consistent signal sequence generation precision above ten percent, which surpasses not only the witness rate as control but the natural rate of binders among unique sequences in the original data (i.e., one percent). In this dataset, in contrast to S_single and S_poly, there are signal-unrelated systematic differences between the repertoires. This can be seen by the distribution of signal-agnostic topics: in S_poly and S_single, all repertoires have identical proportions of these topics (Fig. S1 and Fig. S2, middle columns), as there are no systematic differences except the signal itself, while in SE_absolut, the proportion of each signal-agnostic topic varied across repertoires (Fig. S5, middle column). In this section, we showed that AIRRTM captures complex, 3D-structure-defined synthetic signal – Absolut!-computed (Robert et al. 2022) binding to a given antigen – in experimental data, and consistently generates novel sequences classified as binders by Absolut!.

### AIRRTM in the wild: repertoire classification and estimation of witness rate for experimental data without ground truth labels for individual sequences

To test AIRRTM in the most challenging and biotechnology-relevant environment, we applied it to the E1 dataset (Gidoni et al. 2019, see Methods 1) containing naïve B cell repertoires from 50 celiac-diseased individuals and 46 healthy controls. In the absence of ground truth labels for individual sequences, we validated our model by classifying unseen repertoires by disease label (celiac, healthy control) using predicted signal intensity levels. We used 5-fold cross-validation, thus training AIRRTM on 80% of the repertoires each time. We trained and used the model to predict signal intensity for sequences of all repertoires. Since we assumed that only a small fraction of sequences define the repertoire label, we represented each repertoire by the lowest signal intensity value of its top 0.01%, 0.02%, … sequences, which are the quantile signal intensity thresholds of the top sequences, as predictors of its label (Fig. 4A). We represented each repertoire as a vector of such quantile thresholds (`q` ∊ `[90%, 99%, 99.5%, 99.8%, 99.9%, 99.95%, 99.98%, 99.99%]`), and trained a simple decision tree classifier on these vectors. Then, we used it to predict labels for the repertoires from the testing set. This method yielded an F1 score of 0.74±0.029 (mean±standard error was obtained by 5-fold cross-validation, while the F1 score for shuffled labels was 0.51±0.048 Fig. 4B), thus matching the performance shown by Shemesh et al. 2021.

**Figure 4:**
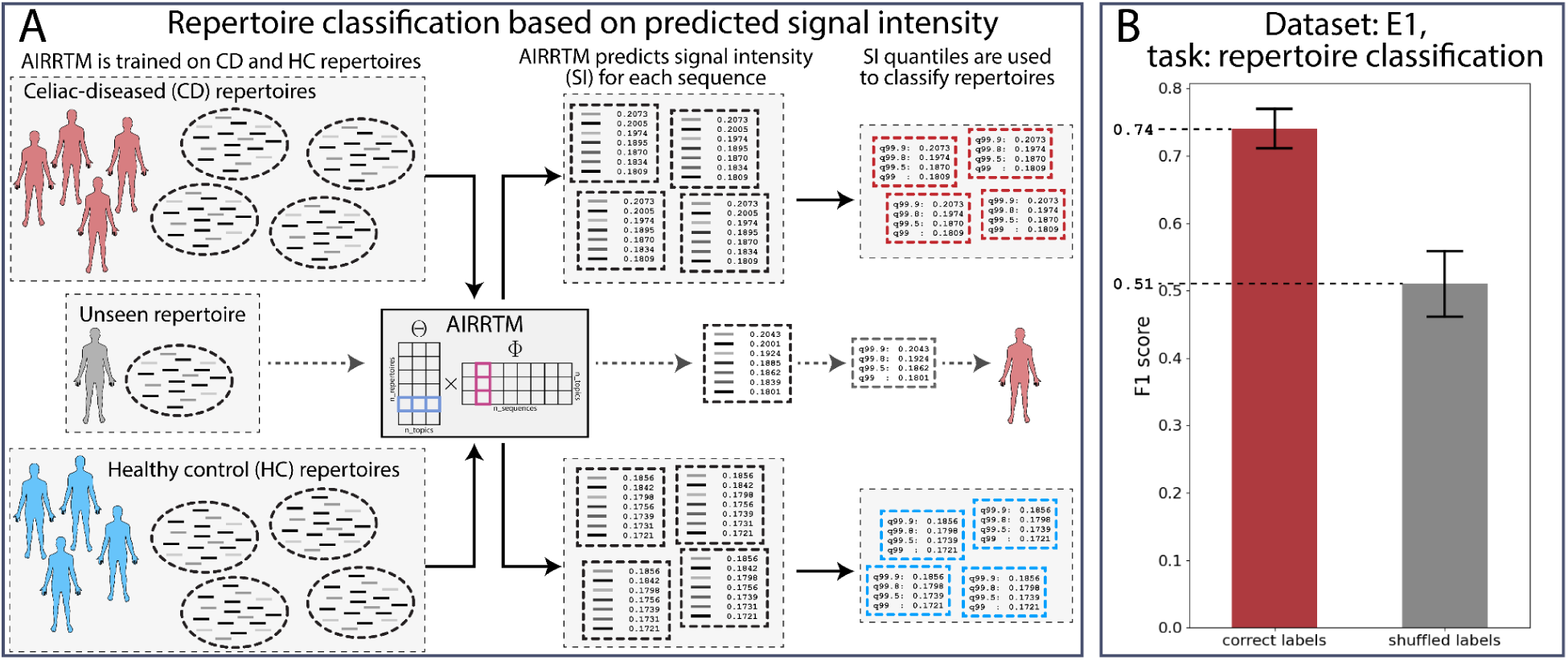
AIRRTM stratifies fully experimental repertoires from Celiac diseased and healthy individuals. (A) Classification of unseen repertoires with AIRRTM: first, the model is trained on the repertoires with known labels – in this case, 50 Celiac-diseased (CD) and 46 Healthy Control (HC) individuals (dataset E1). The trained model is then used to predict signal intensity for all sequences in each repertoire. Each repertoire is represented as a vector of quantiles of predicted signal intensities of its sequences. The repertoires are classified based on the computed quantiles. (B) F1 score for repertoire classification obtained with 5-fold cross-validation of the E1 dataset for correct (red) and shuffled (gray) repertoire labels. The bar height signifies the mean value, the error bars reflect the full range of values obtained by 5-fold cross-validation.

Using predictions for individual sequences to classify repertoires allows us to (i) roughly estimate the true witness rate in the data: if, for example, `q=99.5%` is a strong predictor for repertoire classification, we can assume that the true witness rate of signal-associated sequences is close to or lower than `0.5%`; and (ii) report the top `1-q` (e.g., `0.5%`) sequences ranked by predicted signal intensity as candidates for being signal-associated. Since AIRRTM does not always reach perfect sequence identification precision, some of the reported sequences may be false positives, so `0.5%` in our example is actually an upper bound on the witness rate. For the E1 dataset, `q=99.5%` was indeed a strong predictor. We do not report the top `0.5%` sequences, as the sample labels are not publicly available, and these sequences may be used to reconstruct the labels.

To summarize, we showed that AIRRTM can capture the signal from fully experimental data, and in addition, we showed how it can be used to estimate the true witness rate in the data.

## Discussion

We developed AIRRTM, which identifies immune signal-associated AIR sequences and generates novel ones using repertoire labels without requiring sequence labels. First, we formalized this problem as two ML tasks (weakly supervised signal sequence identification and signal sequence generation), and then implemented an ML approach (AIRRTM) combining the latent representation capacity of sequences from a Sequence VAE and the distribution of sequence usage inside repertoires using Topic Modeling. We validated AIRRTM performance on several increasingly complex datasets: synthetic data with single immune signal, synthetic data with multiple immune signals, experimental data synthetically annotated with a very complex immune signal, and fully experimental data. For each dataset, except the experimental one, ground truth was available and we tested AIRRM performance on different witness rates and showed that in each case it achieves performance sufficient for potential further experimental validation of identified and generated signal-positive sequences.

An ideal use case scenario for AIRRTM would be to train it on a dataset of N repertoires from individuals labeled with an immune-mediated or infectious disease (e.g., Celiac disease or HIV) and N repertoires from healthy controls, identify the sequences that determine the repertoire labels in the positive repertoires and generate novel sequences, and, finally, test these sequences experimentally against the known antigens, associated with the disease, or to use identified and generated sequences as a large set of biomarkers of disease state. We approximated this scenario *in silico* (Fig. 3) with antigen-binding annotated repertoires using Absolut! (Robert et al. 2022). Note that for this scenario to be successful, one does not need sequence identification and generation precision close to 100% – a lower but consistent number (e.g., 10%) would already be of therapeutic interest for high sensitivity. This way, using AIRRTM, one can approach antibody design by leveraging large-scale databases where only repertoire labels exist (Olsen et al. 2022a; Corrie et al. 2018).

An important consideration for AIRRTM is the fact that disease-associated sequences do not necessarily have to be disease-related antigen binders: for example, CMV-associated sequences identified by Emerson et al. 2017 do not match known CMV binders. Such sequences could be binders to unknown epitopes or unknown antigens, or even just be correlated with the actual binders. Nevertheless, within the field of candidate therapeutics discovery, identification of immune-status associated clones from immune-states labeled repertoires without explicit receptor-based antigen labeling is common practice (Laustsen et al. 2021; Parameswaran et al. 2013; Lee et al. 2014; Parola et al. 2018; Brüggemann et al. 1989). Therefore, *in silico* identification and generation of disease-associated candidate sequences using only repertoire-level AIRR data may significantly enhance the speed of systems immunology-driven drug discovery.

We believe that this study, together with Pradier et al. 2023, opens a number of new paths in the generative modeling of AIR repertoires. Recently, many modern machine learning methods have resulted in building foundation models (Ouyang et al. 2022; Jumper et al. 2021; Rives et al. 2021; Kirillov et al. 2023; Madani et al. 2020; Vu et al. 2022, 2023) – very large models that can be quantized and/or used for zero-shot learning or fine-tuned to a certain task. A crucial benefit of foundation models is that they significantly lower the ‘entry threshold’ – the necessary level of resources for building an ML application. Some notable examples are large language model-powered retrieval-augmented chatbots (e.g., [Kindly AI](https://www.kindly.ai/)), generation of illustrated stories (e.g., [Novel AI](https://novelai.net/)), or automatic image annotation (e.g., [Roboflow](https://roboflow.com/)). We argue that foundation models in biology and in AIRR research in particular, may be of even greater benefit, lowering the bar for researchers with less computer science and machine learning background, similarly to how immuneML (Pavlović et al. 2021) lowers this bar. We hypothesize that using a Large Protein Language Model such as ProtGPT2 (Ferruz et al. 2022), ProtTrans (Elnaggar et al. 2021), or ProGen2 (Nijkamp et al. 2022) as a foundation model for AIRRTM encoder may lead to (i) significantly improving AIRRTM performance and (ii) lead AIRRTM to become a foundation model for AIR repertoires.

To conclude, in this work, we showed that immune signals can be inferred from data where only repertoire labels are available and that the inferred signals can be used to generate novel signal sequences – thus broadening the search space for AIR-based therapeutics by allowing for the discovery of drug candidates in large-scale publicly available experimental data (Corrie et al. 2018; Olsen et al. 2022a).

## Methods

### Methods 1. Definition of immune signal in synthetic and experimental immunoglobulin sequencing data

We define a *generic immune signal* (Pavlović et al. 2021) as a binary function defined on the set of all AIR sequences (of length<N, to make this set finite). In other words, an immune signal is a property of an AIR sequence that can be present or not in any given sequence. Each immune signal has its *natural occurrence rate* in a given repertoire: the probability of observing a signal-positive sequence in this repertoire.

Thus, any set of AIR sequences can define an immune signal. In this study, however, we focus on several types of simpler immune signals that can be defined with specific rules and easily identified as ground truth. The most primitive one is the presence of a given (gapped) k-mer. Using it as a starting point, one can construct more complex immune signals, such as “two different k-mers are present at the same time” (which can be considered a gapped k-mer with varying length) or “at least one of two different k-mers is present”. The first rule is more restrictive with respect to the original rule (i.e., a single k-mer) because it is constructed via intersecting two immune signals, while the second one is less restrictive since it is constructed as a union of two immune signals. Using a too restrictive rule for defining an immune signal may lead to a too low natural occurrence rate of this signal in the data which, in turn, makes it infeasible to generate natural-like datasets containing this immune signal, i.e., the natural occurrence rate of this signal is too low (see Fig. S3 for examples of natural occurrence rate). We therefore focused on less restrictive rules such as sequence motifs: the S_poly dataset is constructed this way: the signal there is “at least one of the four signal amino acid 3-mers is present”.

As the main goal of this study was to analyze the identifiability and reproducibility of immune signals in different settings, we used several synthetic datasets of increasing complexity. All datasets were generated using IGoR (Marcou et al. 2018) and rejection sampling for immune signal-positive sequences: first, with fixed RGMPs (Slabodkin et al. 2021), a large pool of sequences is generated and separated into pools of immune signal-positive and -negative sequences. Then, to generate a sample of size N with *α*⋅N immune signal-positive sequences (with a witness rate *α*), we sample immune signal-positive and -negative sequences from the corresponding pools. Note that we used IGoR to generate full-length heavy-chain antibody sequences and identified CDRH3 using MiXCR (Bolotin et al. 2015).

For all datasets, we generated 100 samples of 80000 sequences each (repeated for witness rate in [0.01%, 0.02%, 0.05%, 0.1%, 0.2%, 0.05%]):

- S_single: 50 immune signal-positive, 50 -negative samples. All samples generated with the same RGMPs (RGMP_1). Immune signal is a single amino acid 3-mer.
- S_poly: 50 immune signal-positive, 50 -negative samples. All samples generated with the same RGMPs (RGMP_1). Immune signal is a combination of four different 3-mers with varying natural occurrence rate (Fig. S3B-E). The individual 3-mer rates varied across signal-positive repertoires: for each repertoire, we sampled signal proportions from the uniform(0,1) distribution and rescaled them so that they add up to the total witness rate, thus diluting the signal between the 3-mers and rendering the task more challenging.
- SE_absolut: semi-experimental data: 19 naïve B cell samples from Greiff et al. 2017, annotated with specificity to 140 antigens (Fig. 3A) using the Absolut! software package (Robert et al. 2022). Then, we ranked the samples by the amount of binders to a given antigen, and stripped the bottom half of any binders, thus obtaining immune signal-negative samples. Note that we used an antigen with a natural occurrence rate higher than the maximum witness rate, so we did not have to increase the number of binders.

In addition to these synthetic, and semi-experimental datasets, we also used experimental data from Gidoni et al. 2019:

- E1: 96 naïve B cell samples – 50 from celiac-diseased subjects and 46 from healthy controls. In this dataset, there are no ground truth labels for individual sequences – only repertoire labels.

In all datasets, we used amino acid IGH CDR3 sequences.

### Methods 2. An overview of generative modeling in AIR repertoire research

In machine learning, a generative model is a model that considers the joint probability distribution `P(X,Y)` on given observable variable `X` and target variable `Y`, as opposed to a discriminative model that considers the conditional probability `P(Y` ∣ `X=x)` of the target `Y`, given an observation `x` (Ng and Jordan 2002). From the definition of conditional probability, `P(X, Y) = P(Y | X) · P(X)`: thus, generative models differ from discriminative ones by taking into account `P(X)`, i.e., the distribution of the observable variable – the data. The major advantage of generative models is their ability to generate novel, “unseen” data that follow the same distribution that the existing data was sampled from. Combined with modern representation learning techniques, generative models led to major breakthroughs in *in silico* image, text, and even functional protein synthesis (Kingma and Welling 2014; Brown et al. 2020; Madani et al. 2023).

In the field of AIR repertoire research, several methods were created to model the process of AIR sequences generation, such as an explicit Bayesian IGoR (Marcou et al. 2018) and/or deep generative models (Davidsen et al. 2019; Isacchini et al. 2021). Another notable work is AIRIVA (Pradier et al. 2023) – a generative model that learns interpretable and label-specific representations of TCR repertoires. This approach has led to very promising results, being able to reflect systematic differences between repertoires and identify the TCR sequences responsible for these differences. This work by Pradier et al. focused “on discrete, count representation of TCR repertoires, and they did not directly use the amino-acid sequence information”. The idea of treating each TCR sequence as a unique, discrete entity, while suitable for T cell repertoires, cannot be applied to B cells in many cases. First, repertoire overlap is lower for B cells, while the diversity is extremely high (Briney et al. 2019; Robins et al. 2010; Ortega et al. 2021). Second, the signal in immunoglobulin sequences often involves only a small part of the sequence, not the whole CDR3 (Akbar et al. 2021; Pavlović et al. 2021; Robert et al. 2022). Therefore, it is important to incorporate amino acid sequence into the model of immunoglobulin sequences. Such an approach would also enable generating novel label-specific sequences, which AIRIVA is not designed to do – the model generates repertoires by sampling TCR sequences from a predefined pool.

In addition to modeling of repertoires, deep generative models were used for *in silico* antibody design (Davidsen et al. 2019; Amimeur et al. 2020; Friedensohn et al. 2020; Widrich et al. 2020; Eguchi et al. 2020; Saka et al. 2021; Shuai et al. 2021; Akbar et al. 2022b). The problem with the latter approach is that while repertoire labels are relatively common (e.g., an AIR repertoire from a SARS-CoV-2-positive individual), sequence labels (e.g., a sequence of an antibody specific to SARS-CoV-2) still remain scarce and obtaining such data requires much more effort (Greiff et al. 2020; Setliff et al. 2019; Mason et al. 2021; Makowski et al. 2022). While successful in the tasks they were designed for, the aforementioned models are not focused on obtaining representations for repertoires. At the same time, there are several discriminative models that achieve great results in both repertoire- and sequence-level classification, and provide representations for both AIR sequences and AIR repertoires (Emerson et al. 2017; Widrich et al. 2020). In this study, we aimed to bridge this gap: build a model that employs repertoire labels to learn semantic representations of AIR sequences and AIR repertoires and to generate novel sequences with predefined labels (e.g., using samples from SARS-CoV-2-positive and SARS-CoV-2-negative individuals, generate SARS-CoV-2-specific sequences without any prior knowledge on the sequence labels).

### Methods 3. Topic Modeling on AIRR-seq data

Topic modeling is a machine learning and natural language processing technique for identifying patterns (“topics”) in a collection of texts that originates from probabilistic Latent Semantic Analysis (Hofmann 1999; Papadimitriou et al. 2000), where each text is represented with relative frequencies of occurrences of all words from the combined vocabulary in it. In brief, a list of topics is defined by their vocabulary usage, and a text is defined by a probabilistic combination of these topics. In this section, we describe the Topic Modeling methods and derive the adaptation of this method to AIR repertoire sequencing data.

In pLSA, a corpus of texts is represented as a Term Frequency matrix (TF): a matrix of size *vocabulary_size*×*n_texts* where TF[i,k] is the relative frequency of the *i*-th word in the vocabulary in the *k*-th document. Each text, hereby, is represented as a column in this matrix. Then, a very unrealistic assumption is made that each text can be generated in the following way: there are *n_topics* topics, and, repeatedly, for a given text (i) a topic is drawn from a discrete distribution over all topics that is specific to this text (ii) a word is drawn from a distribution over all words in the vocabulary that is specific to this topic. This way, if we build a matrix Θ of size *n_topics*×*n_texts*, where each column Θ[:,k] represents the topic proportions of the *k*-th text; and a matrix Φ of size *vocabulary_size*×*n_topics*, where each column Φ[:,j] represents the words distribution of the *j*-th topic, the product Φ×Θ will give us a matrix M that is closely related to the TF matrix. The size of M is *vocabulary_size*×*n_texts*, and M[i,k] = Σ*_j_* Φ[i,j]Θ[j,k] is the probability of observing the *i*-th word from the vocabulary in the *k*-th document, and the TF matrix represents the likelihood of our data. In other words, Φ and Θ should be optimized to minimize the difference between Φ×Θ and TF. The described model is generative – one can use it to generate new texts with term frequencies that resemble the texts from the corpus – however, the goal here is to learn *representations*: the columns of Φ represent words and can be used as word embeddings, while the columns of Θ represent documents and can be used as document embeddings. Even topics, despite being constructed implicitly, can be inspected: pLSA is capable of producing interpretable topics: a topic is characterized by a few words that have the highest probability in its distribution. E.g., these few words may be related to football, politics, or adaptive immunity.

A number of extensions of pLSA were created; the most notable are: Latent Dirichlet Allocation (Blei et al. 2001) – by far the most widely used method in Topic Modeling, which introduces Dirichlet priors for Φ and Θ that helps to reduce overfitting substantially; Additive Regularization of Topic Models (Vorontsov and Potapenko 2015) – a semi probabilistic approach that combines pLSA with flexible multi-purpose regularizers; and Embedded Topic Models (Dieng et al. 2020) that combines pLSA with word embeddings. Kazwini and Sanguinetti 2023 applied an LDA-based model to multi-omic single cell data.

Using Topic Modeling to compare AIR repertoires has both intrinsic advantages and disadvantages. Much unlike words in natural texts, AIR sequences in (naïve) repertoires are generated independently, which makes our main assumption less questionable. On the other hand, using one-hot encodings/explicit embeddings limits the amount of information available to the model, since the diversity of AIR sequences supersedes that of the vocabulary of any human language by many orders of magnitude (though for TCRs, Pradier et al. 2023 indeed used a vocabulary of all TCR sequences in the dataset). To overcome this problem, we used an LSTM-based encoder-decoder architecture that, instead of assigning an initially random latent space vector to each AIR sequence, puts the sequence into a latent space using an LSTM encoder network.

Our proposed AIRRTM model (Fig. S6A) combines the ideas of ETM, ARTM and LSTM encoder-decoder in the following manner:

- Given a set of repertoires {R_k_}, where each repertoire is a set of sequences R_k_ ={s_i_}
- The model architecture consists of:

○ A LSTM-based VAE:

▪ A LSTM-based encoder network Enc that transforms a nucleotide AIR sequence into a pair of L-dimensional vectors: *μ* and *σ*, where L is a hyperparameter of the model – the latent space size.
▪ A LSTM-based decoder network Dec that transforms a L-dimensional vector sampled from the normal distribution N(*μ*,*σ*) with mean=*μ* and diagonal covariance matrix Diag(*σ*) back to a nucleotide sequence (with a softmax recurrently returning a probability for each nucleotide to be generated).
○ The topic proportions usage of each repertoire, as a matrix Θ of size `n_topics×n_repertoires`, where each column Θ`[:,k]` represents the topic proportions of the `k`-th repertoire. The values of Θ are parameters that are also optimized directly during training (model weights).
○ A sequence probability matrix Φ of size `n_sequences×n_topics`, where each column Φ`[:,j]` represents the sequence probabilities of the *j*-th topic. The values of Φ are not optimized during training but instead are derived from Enc: Φ[i,j] = σ(Encoder(s_j_)) – i.e., by the cosine similarity between the vector of encoded sequence and the topic embedding

▪ in fact, we do not keep Φ explicitly anywhere, we only use its values during training and inference, similar to the processing of sparse matrices in collaborative filtering (Rendle 2010).
○ A loss function, that itself consists of three additive terms (with weights that are hyperparameters of the models):

▪ the TM likelihood: P(s|R_k_) = Σ*_j_*P(s|t_j_)·P(t_j_|R_k_) = Σ*_j_* σ(Enc(s)) · Θ[j,k] – the likelihood of sequence s originating from repertoire R_k_.
▪ the VAE loss:

- the reconstruction loss: probability (mean cross-entropy for all symbols of the sequences as predicted by the decoder softmax).
- The Kulback-Leibler divergence between the Normal distribution defined by the encoded sequence and the Standard Normal distribution.
▪ Binary cross entropy of repertoire classification for the logistic regression on repertoire topic proportions.

Thus, our model can be considered a Sequence-VAE, where the directions in the latent space represent topics in the topic model. Although we used LSTM for full-length modeling or relatively short (∼20 symbols) sequences, one could as well use another sequential architecture (e.g., transformer or stacked convolutions). Note that if one dropped the generative part of the model (decoder and repertoire topic proportions) and used the latent representation of a sequence to predict repertoire label, the setting becomes identical to noisy-label-learning formulated by Chen et al. 2023.

Once the model is trained, we use the learned repertoire topic proportions to predict signal intensity (SI) for sequences: first, we compute topic weights – for each topic, the difference between its average proportion in positive and negative repertoires; and normalize the vector of weights to have L_1_-norm `= 1`. The signal intensity of a sequence is then calculated as a dot product of its predicted topic probabilities vector and the normalized topic weights vector.

### Methods 4. Detailed architecture of the AIRRTM encoder

We trained an AIRRTM model for two objectives: detect an immune signal in the AIR sequences and learn latent space representations for them. In terms of the loss function, the first objective consists of two parts: the classical Topic Modeling loss (i.e., predict if a sequence S comes from a repertoire R by using topic proportions for R and predicting topic probabilities for S) and the repertoire classification loss (i.e., predict the repertoire label using its topic proportions). The second objective is defined by the VAE loss, which consists of the sequence reconstruction loss and the Kullback-Leibler divergence. These two objectives may differ in the nature of what they need to learn: for example, the immune signal may involve only a small part of the receptor, and thus for predicting if a sequence originated from a signal-positive or a signal-negative repertoire (objective 1) the model needs access to local sequential features. At the same time, to reconstruct the sequence (objective 2) one needs to process its full length. However, we cannot just employ two separate models because to generate signal-related sequences successfully, we require the sequence latent space representations to represent the immune signal semantically. We overcome this apparent contradiction by two steps.

The first step is using a two-component encoder (Fig. S6B): we pass the input through an LSTM recurrent network and a more simple convolutional network with skip connections. The LSTM (accompanied by an LSTM decoder) handles the sequence reconstruction. At the same time, the second component detects the signal and can be tailored to detect the signal the user expects to observe in the data (for example, if we know that the signal is a set of k-mers, we can use a single 1D-convolutional layer with multiple filters, or we can incorporate computation of physicochemical properties if we suspect the signal to have a physicochemical nature). The outputs of the two components are then concatenated and passed through a single fully connected layer to obtain the latent-space representation. The second step is not simply using the axes of the latent space as topics but instead adding another linear layer without activation function that projects latent_space_size (large) dimensions into n_topics (low/task-dependent) dimensions. This way, the topics are directions in the latent space, and we can utilize it to sample the signal-related sequences from these topics.

### Methods 5. Sampling from signal-related topics

The encoder and the VAE sampling step transform an AIR sequence into a distribution in the latent space. If the VAE is trained correctly, the latent space is continuous (cite Kingma and Welling 2013), meaning that if we take a (reasonably) arbitrary point in the latent space and pass it through the decoder, we will get a sequence that could be observed in the data. By reasonably, we mean that the KL-divergence in the loss keeps all sequence representations close to zero, so if we take a too-distant point, its decoding will likely result in a non-AIR-like sequence.

Recall that topics are directions in the latent space. Ideally, to generate signal-positive AIR sequences, if we found an immune signal-related topic, we would want to take a point that is located very far along this direction. However, the concentration of the data embeddings around zero does not allow us to do that. Another solution is to take the signal-positive AIR sequences identified in the data, calculate their mean and standard deviation, and then sample from the area these sequences cover in the latent space – i.e., sample from the normal distribution with the calculated mean and standard deviation (Fig. S6C). We chose the latter solution as the one yielding more natural-like sequences.

## Software

We used IGoR v1.4.0 (Marcou et al. 2018) for sequence generation and MiXCR v4.4.1 (Bolotin et al. 2015) for sequence annotation.

## Graphics

Matplotlib v3.3.2 (Hunter 2007). Seaborn v0.11.0 (Michael Waskom et al. 2020).

## Hardware

Computations were performed on a dedicated server as well as the high-performance computing cluster FRAM (Norwegian e-infrastructure for Research and Education sigma2.no/fram).

## Software Availability

AIRRTM is available in a Github repository, https://github.com/csi-greifflab/airrtm.

## Data Availability

## Acknowledgments

## Acknowledgements

The authors would like to thank Maria Chernigovskaya, Lonneke Scheffer, and Milena Pavlović for discussions on the nature of immune signals.

## Supplemental Materials

**Figure S1 (relates to Fig. 2):**
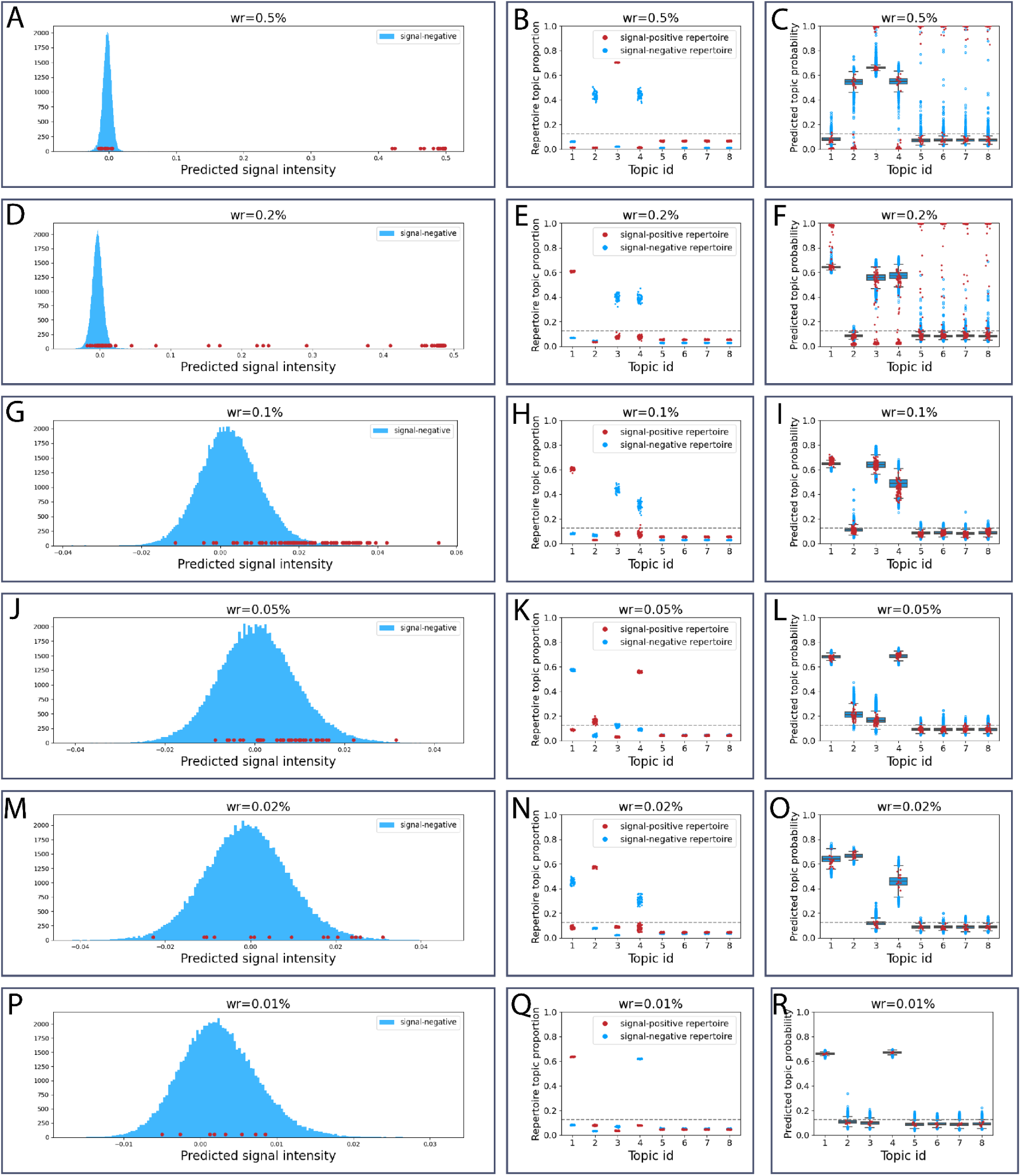
Per-sequence signal strength predictions and repertoire topic proportions show that AIRRTM captures the signal even for the lowest witness rate in the S_single dataset. Left column (A, D, G, J, M, P): distribution of predicted signal strength for signal-negative (blue) and signal-positive (red) sequences. Middle column (B, E, H, K, N, Q): topic proportions for signal-negative (blue) and signal-positive (colored by k-mer) repertoires. Each column corresponds to a single topic, each point represents the proportions of the given topic in a repertoire. The dashed line corresponds to 1/n_topics, i.e., neutral topic value. Right column (C, F, I, L, O, R): topic proportions for signal-negative (blue) and signal-positive (red) repertoires. Each column corresponds to a single topic, each point represents the proportions of the given topic in a repertoire.

**Figure S2 (relates to Fig. 2):**
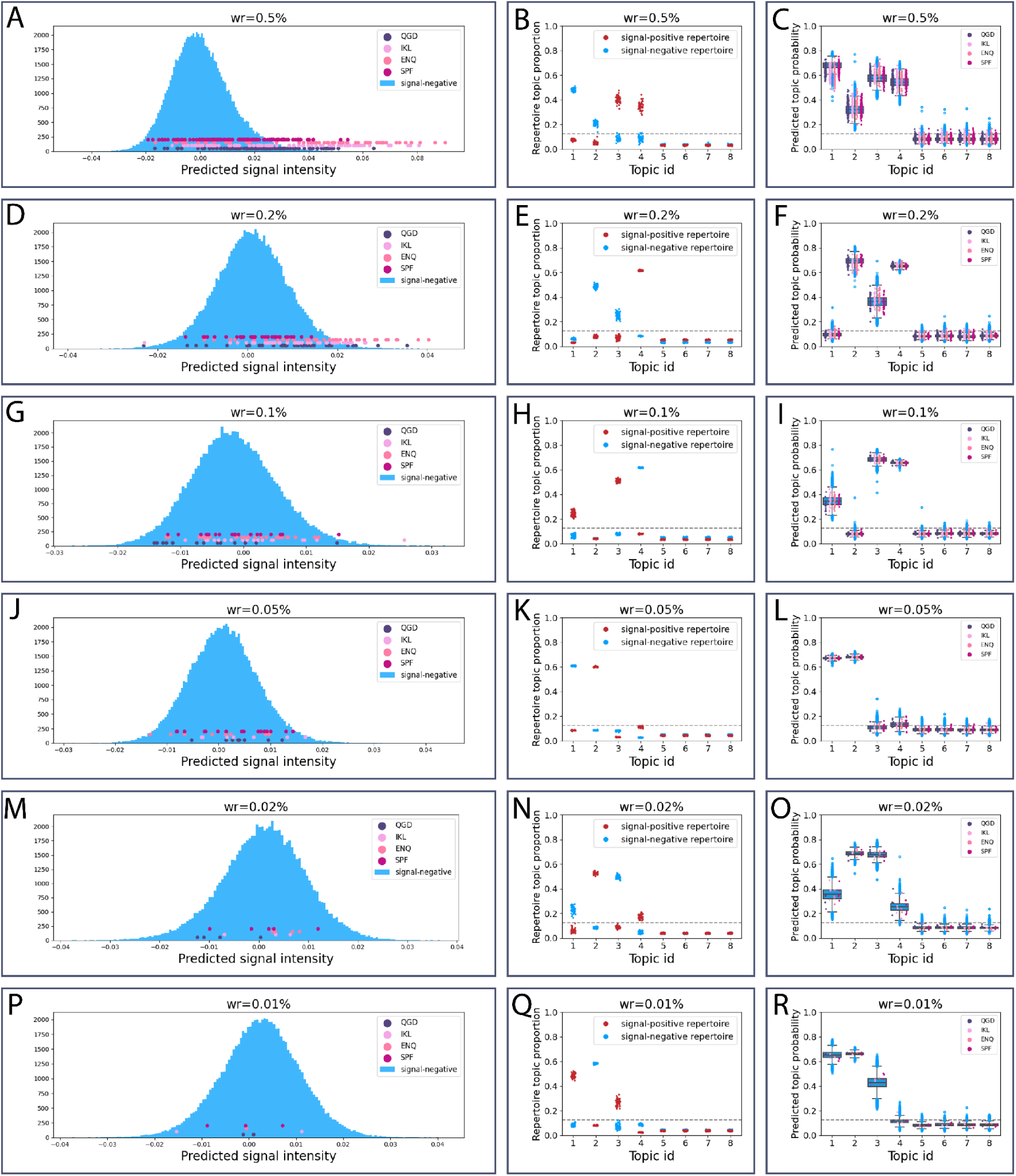
Per-sequence signal strength predictions and repertoire topic proportions show that AIRRTM captures the signal even for the lowest witness rate in the S_poly dataset. Left column (A, D, G, J, M, P): distribution of predicted signal strength for signal-negative (blue) and signal-positive (shades of red, each color corresponds to a single 3-mer) sequences. Middle column (B, E, H, K, N, Q): topic proportions for signal-negative (blue) and signal-positive (colored by k-mer) repertoires. Each column corresponds to a single topic, each point represents the proportions of the given topic in a repertoire. The dashed line corresponds to 1/n_topics, i.e., neutral topic value. Right column (C, F, I, L, O, R): topic probabilities for signal-negative (blue) and signal-positive (colored by k-mer) sequences.

**Figure S3 (relates to Fig. 2):**
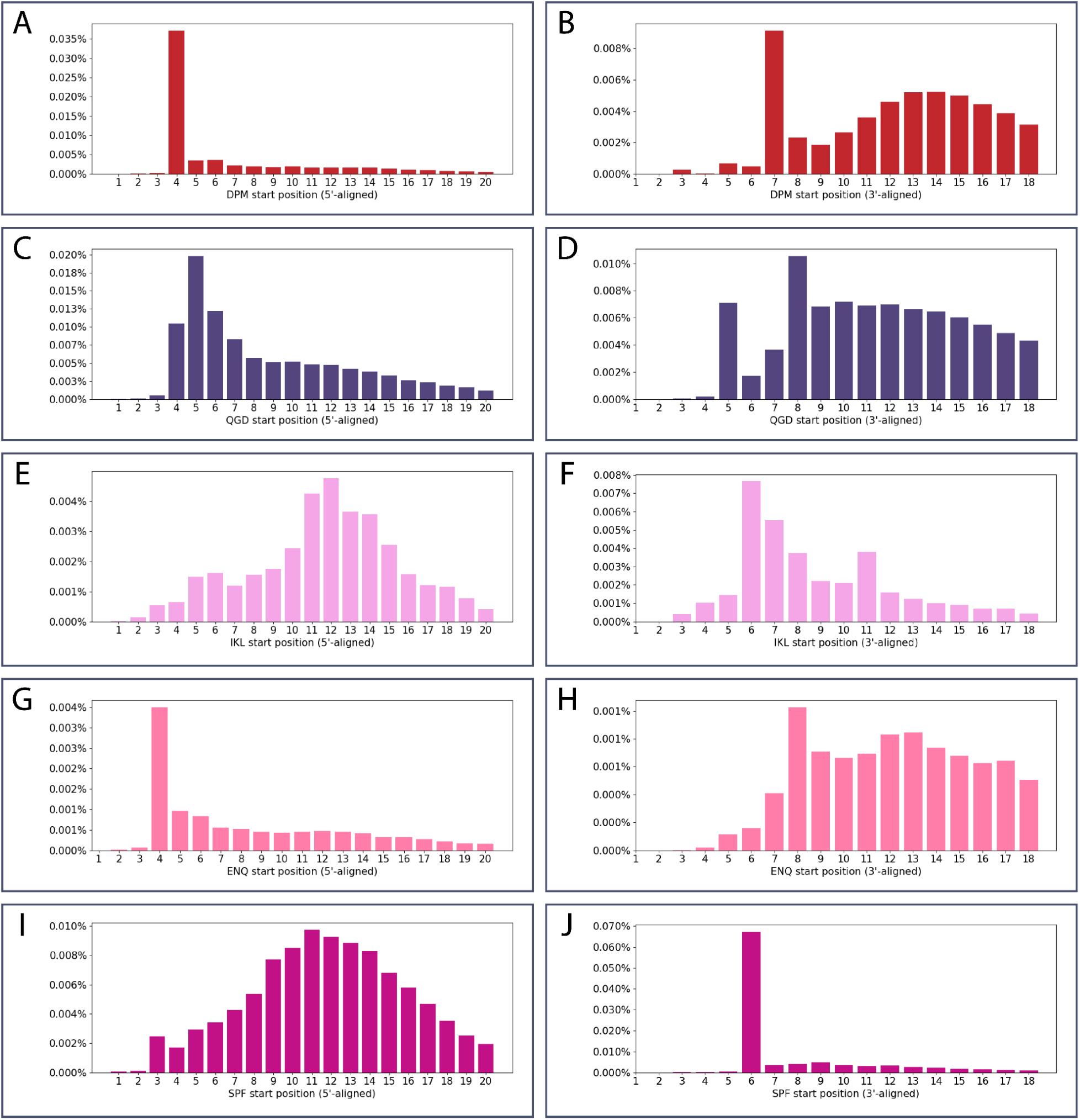
Natural occurrence rate (among IGoR-generated sequences) at different positions for amino acid k-mers representing the signal: k-mer `DKM` in S_single (A–B) and signal k-mers QGD, IKL, ENQ, SPF S_poly (C–J) datasets. The k-mer positions are calculated from the start of the CDR3 (i.e., 5’-aligned, left column) and from the end of the CDR3 (i.e., 3’-aligned, right column).

**Figure S4 (relates to Fig. 2, Fig. 3):**
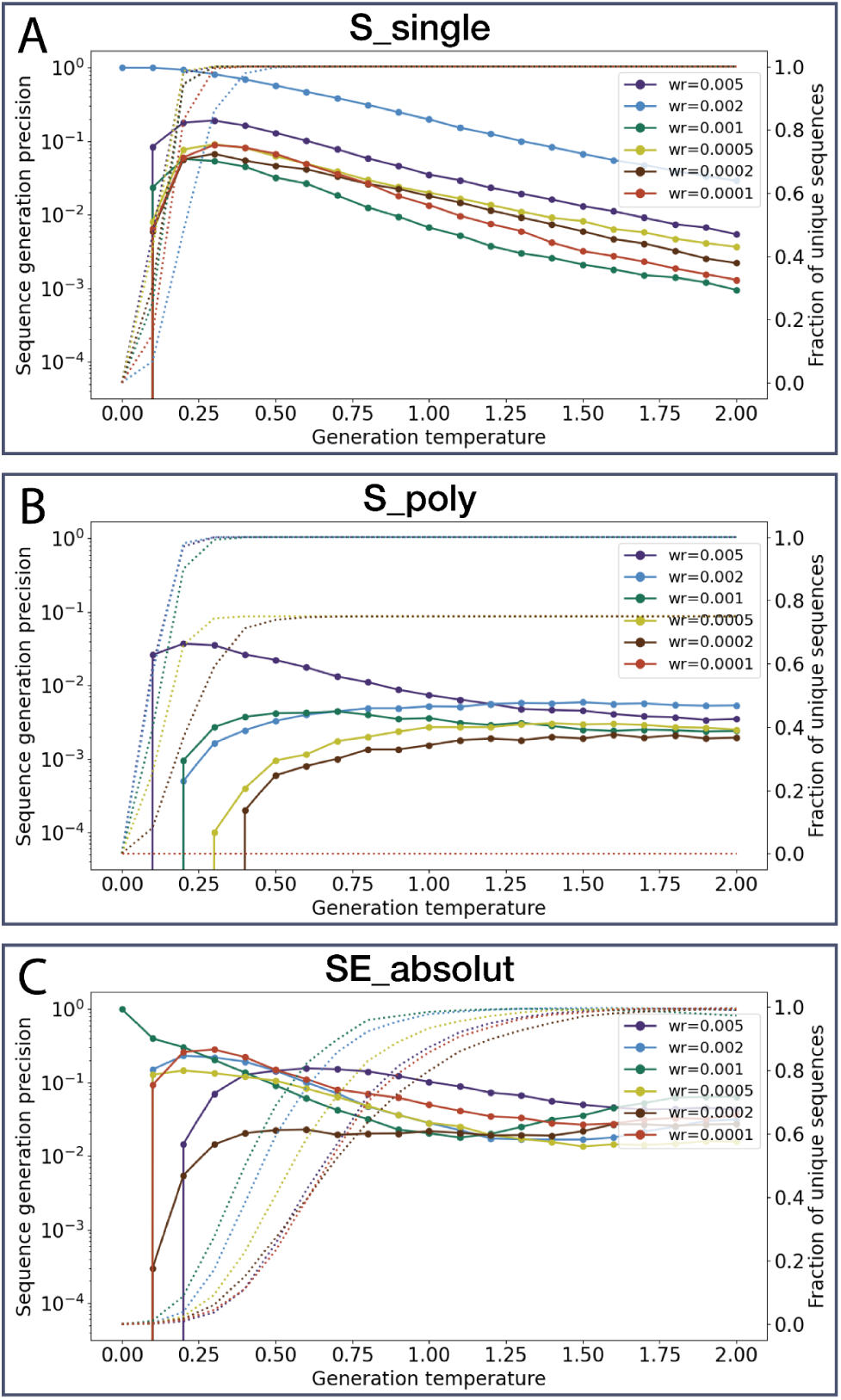
Signal sequence generation precision and diversity as functions of generation temperature. (A) Dataset S_single. Solid lines (Y axis, left) stand for the median generation precision across samples. Dashed lines stand for the fraction of unique sequences of all generated ones. (B) Same as A but for the dataset S_poly. (C) Same as A but for the dataset SE_absolut.

**Figure S5 (relates to Fig. 3):**
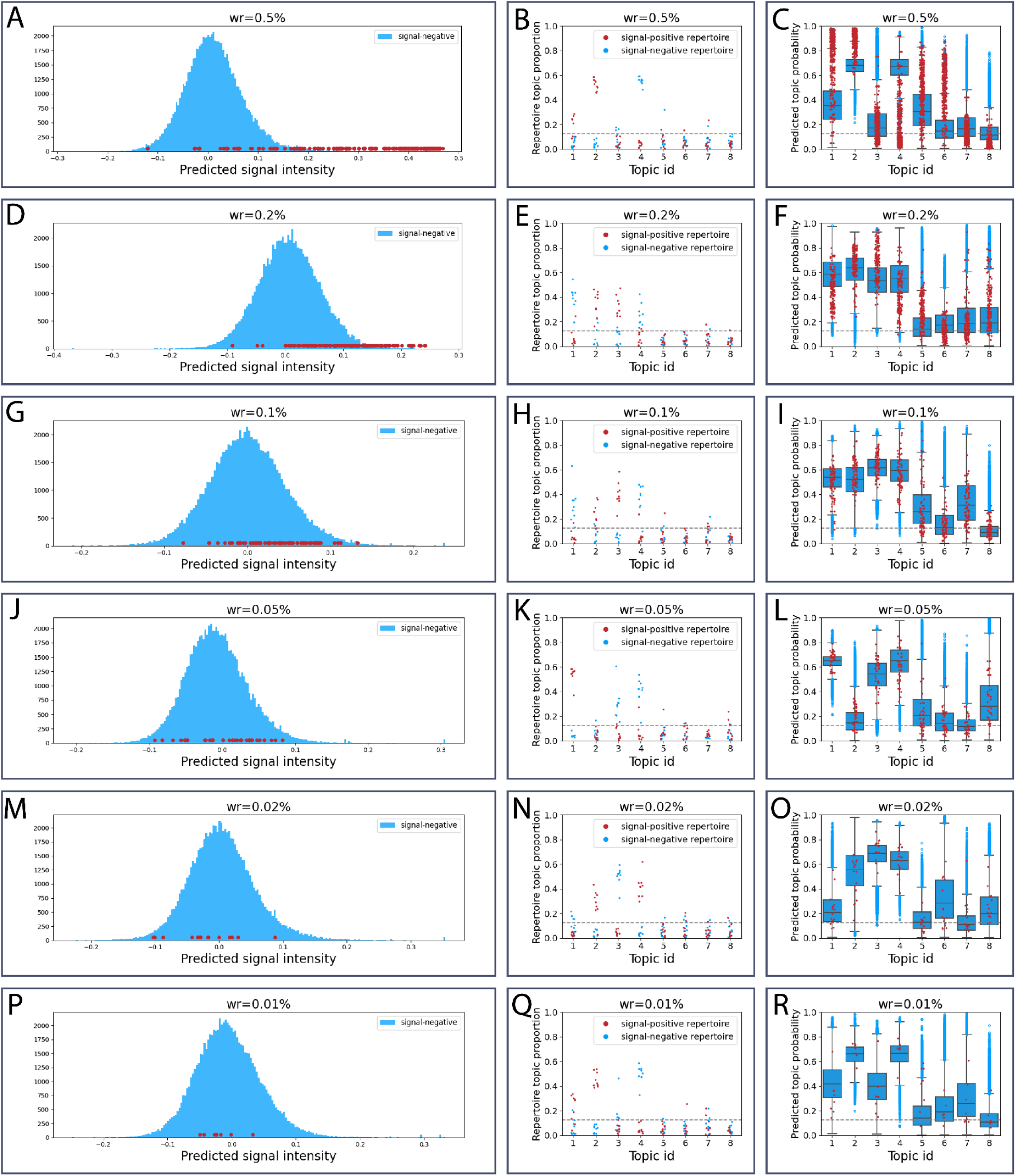
Per-sequence signal strength predictions and repertoire topic proportions show that AIRRTM captures the signal even for the lowest witness rate in the SE_absolut dataset (‘fixed’ scenario). Left column Left column (A, D, G, J, M, P): distribution of predicted signal strength for signal-negative (blue) and signal-positive (shades of red, each color corresponds to a single 3-mer) sequences. Middle column (B, E, H, K, N, Q): topic proportions for signal-negative (blue) and signal-positive (red) repertoires. Each column corresponds to a single topic, each point represents the proportions of the given topic in a repertoire. The dashed line corresponds to 1/n_topics, i.e., neutral topic value. Right column (C, F, I, L, O, R): topic probabilities for signal-negative (blue) and signal-positive (red) sequences.

**Figure S6.**
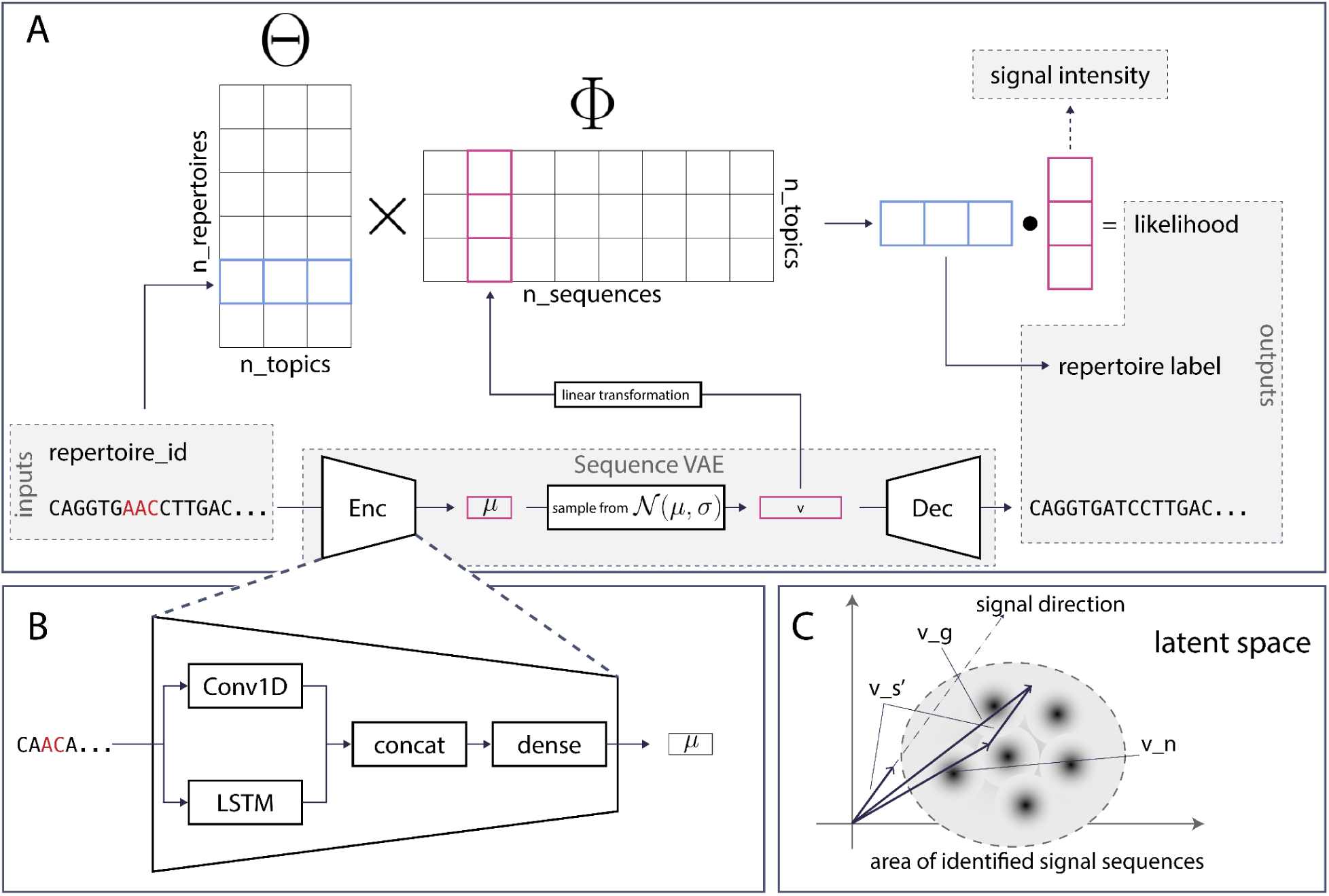
Details of the AIRRTM model. (A) Architecture of the model: (i) a pair (AIR sequence, repertoire_id) is taken as input, (ii) the sequence is encoded to obtain a latent space representation (encoding), (iii) a linear operation with sigmoid transforms the obtained encoding vector into sequence topic probabilities, (iv) using a lookup by repertoire_id, a repertoire topic proportion vector is obtained, (v) the dot product of sequence topic probabilities and repertoire topic proportions is computed as the likelihood of observing the given sequence in the given repertoire – output 1, used to compute the TM loss component, (vi) a repertoire label is predicted by logistic regression on the repertoire topic proportion vector – output 2, used label prediction loss component, (vii) the sequence latent vector is decoded – output 3, used to compute the reconstruction loss component. (B) Architecture of the encoder: an LSTM component responsible for encoding the whole sequence and a convolutional component responsible for capturing the signal. The outputs of both components are concatenated and passed through a single dense layer. (C) Generation of signal-positive sequences by sampling from the area of identified signal-positive sequences.

## References

Abbott RK, Lee JH, Menis S, Skog P, Rossi M, Ota T, Kulp DW, Bhullar D, Kalyuzhniy O, Havenar-Daughton C, et al. 2018. Precursor Frequency and Affinity Determine B Cell Competitive Fitness in Germinal Centers, Tested with Germline-Targeting HIV Vaccine Immunogens. Immunity 48: 133–146.e6.

Akbar R, Bashour H, Rawat P, Robert PA, Smorodina E, Cotet T-S, Flem-Karlsen K, Frank R, Mehta BB, Vu MH, et al. 2022a. Progress and challenges for the machine learning-based design of fit-for-purpose monoclonal antibodies. mAbs 14: 2008790.

Akbar R, Robert PA, Pavlović M, Jeliazkov JR, Snapkov I, Slabodkin A, Weber CR, Scheffer L, Miho E, Haff IH, et al. 2021. A compact vocabulary of paratope-epitope interactions enables predictability of antibody-antigen binding. Cell Rep 34. https://www.cell.com/cell-reports/abstract/S2211-1247(21)00170-4 (Accessed March 24, 2021).

Akbar R, Robert PA, Weber CR, Widrich M, Frank R, Pavlović M, Scheffer L, Chernigovskaya M, Snapkov I, Slabodkin A, et al. 2022b. In silico proof of principle of machine learning-based antibody design at unconstrained scale. mAbs 14: 2031482.

Amimeur T, Shaver JM, Ketchem RR, Taylor JA, Clark RH, Smith J, Citters DV, Siska CC, Smidt P, Sprague M, et al. 2020. Designing Feature-Controlled Humanoid Antibody Discovery Libraries Using Generative Adversarial Networks. 2020.04.12.024844. https://www.biorxiv.org/content/10.1101/2020.04.12.024844v2 (Accessed February 3, 2023).

Blei D, Ng A, Jordan M. 2001. Latent Dirichlet Allocation.

Bolotin DA, Poslavsky S, Mitrophanov I, Shugay M, Mamedov IZ, Putintseva EV, Chudakov DM. 2015. MiXCR: software for comprehensive adaptive immunity profiling. Nat Methods 12: 380–381.

Briney B, Inderbitzin A, Joyce C, Burton DR. 2019. Commonality despite exceptional diversity in the baseline human antibody repertoire. Nature 566: 393–397.

Brown T, Mann B, Ryder N, Subbiah M, Kaplan JD, Dhariwal P, Neelakantan A, Shyam P, Sastry G, Askell A, et al. 2020. Language Models are Few-Shot Learners. In Advances in Neural Information Processing Systems, Vol. 33 of, pp. 1877–1901, Curran Associates, Inc. https://papers.nips.cc/paper/2020/hash/1457c0d6bfcb4967418bfb8ac142f64a-Abstract.html (Accessed February 3, 2023).

Brüggemann M, Caskey HM, Teale C, Waldmann H, Williams GT, Surani MA, Neuberger MS. 1989. A repertoire of monoclonal antibodies with human heavy chains from transgenic mice. Proc Natl Acad Sci U S A 86: 6709–6713.

Chen M, Zhao Y, Wang Z, He B, Yao J. 2023. A Noisy-Label-Learning Formulation for Immune Repertoire Classification and Disease-Associated Immune Receptor Sequence Identification. http://arxiv.org/abs/2307.15934 (Accessed August 6, 2023).

Choi Y. 2022. Artificial intelligence for antibody reading comprehension: AntiBERTa. Patterns 3. https://www.cell.com/patterns/abstract/S2666-3899(22)00132-5 (Accessed September 21, 2023).

Chung J, Kastner K, Dinh L, Goel K, Courville AC, Bengio Y. 2015. A Recurrent Latent Variable Model for Sequential Data. In Advances in Neural Information Processing Systems, Vol. 28 of, Curran Associates, Inc. https://proceedings.neurips.cc/paper/2015/hash/b618c3210e934362ac261db280128c22-Abstract.html (Accessed July 27, 2022).

Corrie BD, Marthandan N, Zimonja B, Jaglale J, Zhou Y, Barr E, Knoetze N, Breden FMW, Christley S, Scott JK, et al. 2018. iReceptor: A platform for querying and analyzing antibody/B-cell and T-cell receptor repertoire data across federated repositories. Immunol Rev 284: 24–41.

Daberdaku S, Ferrari C. 2019. Antibody interface prediction with 3D Zernike descriptors and SVM. Bioinformatics 35: 1870–1876.

Davidsen K, Olson BJ, DeWitt WS III, Feng J, Harkins E, Bradley P, Matsen FA IV. 2019. Deep generative models for T cell receptor protein sequences. eLife 8: e46935.

Dieng AB, Ruiz FJR, Blei DM. 2020. Topic Modeling in Embedding Spaces. Trans Assoc Comput Linguist 8: 439–453.

Eguchi RR, Anand N, Choe CA, Huang P-S. 2020. IG-VAE: Generative Modeling of Immunoglobulin Proteins by Direct 3D Coordinate Generation. bioRxiv 2020.08.07.242347.

Elnaggar A, Heinzinger M, Dallago C, Rehawi G, Wang Y, Jones L, Gibbs T, Feher T, Angerer C, Steinegger M, et al. 2021. ProtTrans: Towards Cracking the Language of Life’s Code Through Self-Supervised Learning. 2020.07.12.199554. https://www.biorxiv.org/content/10.1101/2020.07.12.199554v3 (Accessed August 6, 2023).

Emerson RO, DeWitt WS III, Vignali M, Gravley J, Hu JK, Osborne EJ, Desmarais C, Klinger M, Carlson CS, Hansen JA, et al. 2017. Immunosequencing identifies signatures of cytomegalovirus exposure history and HLA-mediated effects on the T cell repertoire. Nat Genet 49: 659–665.

Ferdous S, Martin ACR. 2018. AbDb: antibody structure database—a database of PDB-derived antibody structures. Database 2018: bay040.

Ferruz N, Schmidt S, Höcker B. 2022. ProtGPT2 is a deep unsupervised language model for protein design. Nat Commun 13: 4348.

Friedensohn S, Neumeier D, Khan TA, Csepregi L, Parola C, Vries ARG de, Erlach L, Mason DM, Reddy ST. 2020. Convergent selection in antibody repertoires is revealed by deep learning. bioRxiv 2020.02.25.965673.

Gao Y, Gao Y, Li W, Wu S, Xing F, Zhou C, Fu S, Chuai G, Chen Q, Zhang H, et al. 2023. Neo-epitope identification by weakly-supervised peptide-TCR binding prediction. 2023.08.02.550128. https://www.biorxiv.org/content/10.1101/2023.08.02.550128v2 (Accessed August 14, 2023).

Gidoni M, Snir O, Peres A, Polak P, Lindeman I, Mikocziova I, Sarna VK, Lundin KEA, Clouser C, Vigneault F, et al. 2019. Mosaic deletion patterns of the human antibody heavy chain gene locus shown by Bayesian haplotyping. Nat Commun 10: 628.

Greiff V, Menzel U, Miho E, Weber C, Riedel R, Cook S, Valai A, Lopes T, Radbruch A, Winkler TH, et al. 2017. Systems Analysis Reveals High Genetic and Antigen-Driven Predetermination of Antibody Repertoires throughout B Cell Development. Cell Rep 19: 1467–1478.

Greiff V, Yaari G, Cowell LG. 2020. Mining adaptive immune receptor repertoires for biological and clinical information using machine learning. Curr Opin Syst Biol 24: 109–119.

Hie BL, Shanker VR, Xu D, Bruun TUJ, Weidenbacher PA, Tang S, Wu W, Pak JE, Kim PS. 2023. Efficient evolution of human antibodies from general protein language models. Nat Biotechnol 1–9.

Hofmann T. 1999. Probabilistic latent semantic indexing. In Proceedings of the 22nd annual international ACM SIGIR conference on Research and development in information retrieval, SIGIR ‘99, pp. 50–57, Association for Computing Machinery, New York, NY, USA 10.1145/312624.312649 (Accessed January 12, 2022).

Hunter JD. 2007. Matplotlib: A 2D graphics environment. Comput Sci Eng 9: 90–95.

Isacchini G, Walczak AM, Mora T, Nourmohammad A. 2021. Deep generative selection models of T and B cell receptor repertoires with soNNia. Proc Natl Acad Sci 118. https://www.pnas.org/content/118/14/e2023141118 (Accessed April 6, 2021).

Jumper J, Evans R, Pritzel A, Green T, Figurnov M, Ronneberger O, Tunyasuvunakool K, Bates R, Žídek A, Potapenko A, et al. 2021. Highly accurate protein structure prediction with AlphaFold. Nature 596: 583–589.

Kanduri C, Pavlović M, Scheffer L, Motwani K, Chernigovskaya M, Greiff V, Sandve GK. 2021. Profiling the baseline performance and limits of machine learning models for adaptive immune receptor repertoire classification. https://www.biorxiv.org/content/10.1101/2021.05.23.445346v2 (Accessed September 29, 2021).

Kazwini NE, Sanguinetti G. 2023. SHARE-Topic: Bayesian Interpretable Modelling of Single-Cell Multi-Omic Data. 2023.02.02.526696. https://www.biorxiv.org/content/10.1101/2023.02.02.526696v1 (Accessed February 24, 2023).

Kingma DP, Welling M. 2014. Auto-Encoding Variational Bayes. arXiv http://arxiv.org/abs/1312.6114 (Accessed May 31, 2022).

Kirillov A, Mintun E, Ravi N, Mao H, Rolland C, Gustafson L, Xiao T, Whitehead S, Berg AC, Lo W-Y, et al. 2023. Segment Anything. http://arxiv.org/abs/2304.02643 (Accessed July 27, 2023).

Kovaltsuk A, Raybould MIJ, Wong WK, Marks C, Kelm S, Snowden J, Trück J, Deane CM. 2020. Structural diversity of B-cell receptor repertoires along the B-cell differentiation axis in humans and mice ed. Y. Ofran. PLOS Comput Biol 16: e1007636.

Laustsen AH, Greiff V, Karatt-Vellatt A, Muyldermans S, Jenkins TP. 2021. Animal Immunization, in Vitro Display Technologies, and Machine Learning for Antibody Discovery. Trends Biotechnol. https://www.sciencedirect.com/science/article/pii/S0167779921000615 (Accessed June 24, 2021).

Lee E-C, Liang Q, Ali H, Bayliss L, Beasley A, Bloomfield-Gerdes T, Bonoli L, Brown R, Campbell J, Carpenter A, et al. 2014. Complete humanization of the mouse immunoglobulin loci enables efficient therapeutic antibody discovery. Nat Biotechnol 32: 356–363.

Lu R-M, Hwang Y-C, Liu I-J, Lee C-C, Tsai H-Z, Li H-J, Wu H-C. 2020. Development of therapeutic antibodies for the treatment of diseases. J Biomed Sci 27: 1.

Madani A, Krause B, Greene ER, Subramanian S, Mohr BP, Holton JM, Olmos JL, Xiong C, Sun ZZ, Socher R, et al. 2023. Large language models generate functional protein sequences across diverse families. Nat Biotechnol 1–8.

Madani A, McCann B, Naik N, Keskar NS, Anand N, Eguchi RR, Huang P-S, Socher R. 2020. ProGen: Language Modeling for Protein Generation. http://arxiv.org/abs/2004.03497 (Accessed July 27, 2023).

Makowski EK, Kinnunen PC, Huang J, Wu L, Smith MD, Wang T, Desai AA, Streu CN, Zhang Y, Zupancic JM, et al. 2022. Co-optimization of therapeutic antibody affinity and specificity using machine learning models that generalize to novel mutational space. Nat Commun 13: 3788.

Marcou Q, Mora T, Walczak AM. 2018. High-throughput immune repertoire analysis with IGoR. Nat Commun 9: 561.

Mason DM, Friedensohn S, Weber CR, Jordi C, Wagner B, Meng SM, Ehling RA, Bonati L, Dahinden J, Gainza P, et al. 2021. Optimization of therapeutic antibodies by predicting antigen specificity from antibody sequence via deep learning. Nat Biomed Eng 5: 600–612.

Michael Waskom, Olga Botvinnik, Maoz Gelbart, Joel Ostblom, Paul Hobson, Saulius Lukauskas, David C Gemperline, Tom Augspurger, Yaroslav Halchenko, Jordi Warmenhoven, et al. 2020. mwaskom/seaborn: v0.11.0 (Sepetmber 2020). https://zenodo.org/record/4019146#.X3xdf1lRUxg (Accessed October 6, 2020).

Ng A, Jordan M. 2002. On Discriminative vs. Generative Classifiers: A comparison of logistic regression and naive Bayes. In Advances in Neural Information Processing Systems, Vol. 14 of, MIT Press https://papers.nips.cc/paper/2001/hash/7b7a53e239400a13bd6be6c91c4f6c4e-Abstract.html (Accessed January 12, 2022).

Nijkamp E, Ruffolo J, Weinstein EN, Naik N, Madani A. 2022. ProGen2: Exploring the Boundaries of Protein Language Models. http://arxiv.org/abs/2206.13517 (Accessed August 9, 2023).

Olsen TH, Boyles F, Deane CM. 2022a. Observed Antibody Space: A diverse database of cleaned, annotated, and translated unpaired and paired antibody sequences. Protein Sci 31: 141–146.

Olsen TH, Moal IH, Deane CM. 2022b. AbLang: an antibody language model for completing antibody sequences. Bioinforma Adv 2: vbac046.

Olson BJ, Matsen FA IV. 2018. The Bayesian optimist’s guide to adaptive immune receptor repertoire analysis. Immunol Rev 284: 148–166.

Ortega MR, Spisak N, Mora T, Walczak AM. 2021. Modeling and predicting the overlap of B- and T-cell receptor repertoires in healthy and SARS-CoV-2 infected individuals. https://www.biorxiv.org/content/10.1101/2021.12.17.473105v1 (Accessed December 21, 2021).

Ostmeyer J, Christley S, Toby IT, Cowell LG. 2019. Biophysicochemical Motifs in T-cell Receptor Sequences Distinguish Repertoires from Tumor-Infiltrating Lymphocyte and Adjacent Healthy Tissue. Cancer Res 79: 1671–1680.

Ouyang L, Wu J, Jiang X, Almeida D, Wainwright CL, Mishkin P, Zhang C, Agarwal S, Slama K, Ray A, et al. 2022. Training language models to follow instructions with human feedback. http://arxiv.org/abs/2203.02155 (Accessed July 27, 2023).

Papadimitriou CH, Raghavan P, Tamaki H, Vempala S. 2000. Latent Semantic Indexing: A Probabilistic Analysis. J Comput Syst Sci 61: 217–235.

Parameswaran P, Liu Y, Roskin KM, Jackson KKL, Dixit VP, Lee J-Y, Artiles KL, Zompi S, Vargas MJ, Simen BB, et al. 2013. Convergent antibody signatures in human dengue. Cell Host Microbe 13: 691–700.

Parola C, Neumeier D, Reddy ST. 2018. Integrating high-throughput screening and sequencing for monoclonal antibody discovery and engineering. Immunology 153: 31–41.

Pavlović M, Scheffer L, Motwani K, Kanduri C, Kompova R, Vazov N, Waagan K, Bernal FLM, Costa AA, Corrie B, et al. 2021. immuneML: an ecosystem for machine learning analysis of adaptive immune receptor repertoires. bioRxiv 2021.03.08.433891.

Pradier MF, Prasad N, Chapfuwa P, Ghalebikesabi S, Ilse M, Woodhouse S, Elyanow R, Zazo J, Gonzalez J, Greissl J, et al. 2023. AIRIVA: A Deep Generative Model of Adaptive Immune Repertoires. http://arxiv.org/abs/2304.13737 (Accessed May 4, 2023).

Rappazzo CG, Fernández-Quintero ML, Mayer A, Wu NC, Greiff V, Guthmiller JJ. 2023. Defining and Studying B Cell Receptor and TCR Interactions. J Immunol Baltim Md 1950 211: 311–322.

Raybould MIJ, Kovaltsuk A, Marks C, Deane CM. 2020. CoV-AbDab: the coronavirus antibody database. Bioinformatics. 10.1093/bioinformatics/btaa739 (Accessed March 10, 2021).

Rendle S. 2010. Factorization Machines. In 2010 IEEE International Conference on Data Mining, pp. 995–1000.

Rives A, Meier J, Sercu T, Goyal S, Lin Z, Liu J, Guo D, Ott M, Zitnick CL, Ma J, et al. 2021. Biological structure and function emerge from scaling unsupervised learning to 250 million protein sequences. Proc Natl Acad Sci 118: e2016239118.

Robert PA, Akbar R, Frank R, Pavlović M, Widrich M, Snapkov I, Chernigovskaya M, Scheffer L, Slabodkin A, Mehta BB, et al. 2021. One billion synthetic 3D-antibody-antigen complexes enable unconstrained machine-learning formalized investigation of antibody specificity prediction. bioRxiv 2021.07.06.451258.

Robert PA, Akbar R, Frank R, Pavlović M, Widrich M, Snapkov I, Slabodkin A, Chernigovskaya M, Scheffer L, Smorodina E, et al. 2022. Unconstrained generation of synthetic antibody–antigen structures to guide machine learning methodology for antibody specificity prediction. Nat Comput Sci 2: 845–865.

Robins HS, Srivastava SK, Campregher PV, Turtle CJ, Andriesen J, Riddell SR, Carlson CS, Warren EH. 2010. Overlap and Effective Size of the Human CD8+ T Cell Receptor Repertoire. Sci Transl Med 2: 47ra64–47ra64.

Ruffolo JA, Gray JJ, Sulam J. 2021. Deciphering antibody affinity maturation with language models and weakly supervised learning. http://arxiv.org/abs/2112.07782 (Accessed September 21, 2023).

Saka K, Kakuzaki T, Metsugi S, Kashiwagi D, Yoshida K, Wada M, Tsunoda H, Teramoto R. 2021. Antibody design using LSTM based deep generative model from phage display library for affinity maturation. Sci Rep 11: 5852.

Sandve GK, Greiff V. 2022. Access to ground truth at unconstrained size makes simulated data as indispensable as experimental data for bioinformatics methods development and benchmarking. Bioinformatics 38: 4994–4996.

Setliff I, Shiakolas AR, Pilewski KA, Murji AA, Mapengo RE, Janowska K, Richardson S, Oosthuysen C, Raju N, Ronsard L, et al. 2019. High-Throughput Mapping of B Cell Receptor Sequences to Antigen Specificity. Cell 179: 1636–1646.e15.

Shemesh O, Polak P, Lundin KEA, Sollid LM, Yaari G. 2021. Machine Learning Analysis of Naïve B-Cell Receptor Repertoires Stratifies Celiac Disease Patients and Controls. Front Immunol 12: 627813.

Shrock EL, Timms RT, Kula T, Mena EL, West AP, Guo R, Lee I-H, Cohen AA, McKay LGA, Bi C, et al. 2023. Germline-encoded amino acid–binding motifs drive immunodominant public antibody responses. Science 380: eadc9498.

Shuai RW, Ruffolo JA, Gray JJ. 2021. Generative Language Modeling for Antibody Design. https://www.biorxiv.org/content/10.1101/2021.12.13.472419v1 (Accessed December 15, 2021).

Slabodkin A, Chernigovskaya M, Mikocziova I, Akbar R, Scheffer L, Pavlović M, Bashour H, Snapkov I, Mehta BB, Weber CR, et al. 2021. Individualized VDJ recombination predisposes the available Ig sequence space. Genome Res 31: 2209–2224.

Stiegler G, Kunert R, Purtscher M, Wolbank S, Voglauer R, Steindl F, Katinger H. 2001. A Potent Cross-Clade Neutralizing Human Monoclonal Antibody against a Novel Epitope on gp41 of Human Immunodeficiency Virus Type 1. AIDS Res Hum Retroviruses 17: 1757–1765.

Swindells MB, Porter CT, Couch M, Hurst J, Abhinandan KR, Nielsen JH, Macindoe G, Hetherington J, Martin ACR. 2017. abYsis: Integrated Antibody Sequence and Structure—Management, Analysis, and Prediction. J Mol Biol 429: 356–364.

Vorontsov K, Potapenko A. 2015. Additive regularization of topic models. Mach Learn 101: 303–323.

Vu MH, Akbar R, Robert PA, Swiatczak B, Sandve GK, Greiff V, Haug DTT. 2023. Linguistically inspired roadmap for building biologically reliable protein language models. Nat Mach Intell 5: 485–496.

Vu MH, Robert PA, Akbar R, Swiatczak B, Sandve GK, Haug DTT, Greiff V. 2022. ImmunoLingo: Linguistics-based formalization of the antibody language. http://arxiv.org/abs/2209.12635 (Accessed July 27, 2023).

Walker LM, Huber M, Doores KJ, Falkowska E, Pejchal R, Julien J-P, Wang S-K, Ramos A, Chan-Hui P-Y, Moyle M, et al. 2011. Broad neutralization coverage of HIV by multiple highly potent antibodies. Nature 477: 466–470.

Weinstein JA, Jiang N, White RA, Fisher DS, Quake SR. 2009. High-throughput sequencing of the zebrafish antibody repertoire. Science 324: 807–810.

Widrich M, Schäfl B, Ramsauer H, Pavlović M, Gruber L, Holzleitner M, Brandstetter J, Sandve GK, Greiff V, Hochreiter S, et al. 2020. Modern Hopfield Networks and Attention for Immune Repertoire Classification. ArXiv200713505 Cs Q-Bio Stat. http://arxiv.org/abs/2007.13505 (Accessed August 14, 2020).

Wu X, Yang Z-Y, Li Y, Hogerkorp C-M, Schief WR, Seaman MS, Zhou T, Schmidt SD, Wu L, Xu L, et al. 2010. Rational Design of Envelope Identifies Broadly Neutralizing Human Monoclonal Antibodies to HIV-1. Science 329: 856–861.

Zhou Z-H. 2018. A brief introduction to weakly supervised learning. Natl Sci Rev 5: 44–53.

